# Membrane voltage dysregulation driven by metabolic dysfunction underlies bactericidal activity of aminoglycosides

**DOI:** 10.1101/2020.04.23.058362

**Authors:** Giancarlo N. Bruni, Joel M. Kralj

## Abstract

Aminoglycosides are broad-spectrum antibiotics whose mechanism of bactericidal activity has been under debate. It is widely accepted, however, that membrane voltage potentiates aminoglycoside activity, which is ascribed to voltage dependent drug uptake. In this paper, we measured the single cell response of *Escherichia coli* treated with aminoglycosides and discovered that the bactericidal action arises not from the downstream effects of voltage dependent drug uptake, but rather directly from dysregulated membrane potential. In the absence of voltage, aminoglycosides are taken into cells and exert bacteriostatic effects by inhibiting translation. However, cell killing was immediate upon re-polarization. The hyperpolarization arose from altered ATP flux, which induced a reversal of the F1Fo-ATPase to hydrolyze ATP and generated the deleterious voltage. Heterologous expression of an ATPase inhibitor from *Salmonella* completely eliminated bactericidal activity, while loss of the F-ATPase significantly reduced the electrophysiological response to aminoglycosides. Our data support a model of voltage induced death, which could be resolved in real-time at the single cell level, and separates the mechanisms of aminoglycoside bacteriostasis and bactericide in *E. coli*.

## Introduction

Aminoglycosides are a potent class of translation inhibitor antibiotics with a broad activity spectrum. Despite a long history in the clinic^1^, their exact mechanism of action remains unclear^2–4^. In gram negative bacteria, aminoglycosides must cross the outer membrane and plasma membrane^5^, into the cytoplasm where they can exert their bactericidal effect which requires binding to the ribosome^6^. The kinetics of uptake into the cytoplasm have been extensively studied and occur in three steps^5^. An ionic interaction between the polycationic aminoglycosides and the outer membrane of the bacterial cell induces a disruption of the outer membrane^7^, and allows the aminoglycoside to ionically associate with the inner membrane^8^. The next step is known as the energy dependent phase I (EDP-I) and occurs almost instantaneously upon aminoglycoside treatment^9^. This portion is noted as energy dependent because both respiration inhibitors^10^ and differential carbon sources^11^ reduced uptake. EDP-I is thought to be the step at which the aminoglycoside enters the cytoplasm^5,11^, is concentration dependent^8^, and occurs in cells that are resistant to or tolerant of aminoglycosides^2,12^. Following EDP-I is EDP-II, which only occurs in aminoglycoside sensitive cells^5,12^, is thought to be essential for the bactericidal activity of aminoglycosides, and requires respiration^12^. Throughout these early studies, uptake of the aminoglycosides was often treated as synonymous with bactericidal activity.

Proposed bactericidal mechanisms all stem from this consensus theory of aminoglycoside uptake^2,4,10,13,14^. Once aminoglycosides are inside the cell, several competing theories exist to explain bactericidal activity including membrane breakdown from mistranslated protein^14,15^, reactive oxygen species^4^ (ROS), and a positive feedback of drug uptake^2,10^, though there is debate around each^2,13,16^. Despite this debate, there is broad agreement upon two important points. The first is that the uptake mechanism, and therefore the resulting bactericidal activity, is voltage dependent^17^. That is, bactericidal activity occurs after uptake, and that uptake is intrinsically tied to membrane potential^2,5,14^. This makes sense given the ample evidence of broken respiration protecting bacteria from aminoglycosides^2,11,18,19^. The second point is that this voltage induced uptake is responsible for mistranslation of protein upon aminoglycoside binding, which in turn creates the membrane breakdown essential for bactericidal activity. These pores, or the ROS produced in their occurrence, are thought to be responsible for the bactericidal activity of aminoglycosides. New techniques offer the ability to study the effects of aminoglycosides and perhaps resolve some debated aspects of its mechanism of action.

Single cell, fluorescent imaging offers a means to shed light on the effects of antibiotic exposure with high resolution in space and time. Improvements in microscope hardware enable automated live cell imaging while resolving the responses of individual bacteria. This hardware can be coupled with genetically encoded, or chemical fluorescent sensors that report bacterial voltage^20–22^, calcium^23^, and ATP^24,25^, providing a lens to explore the long-term effects of antibiotic exposure. Recently, live cell voltage imaging of *Bacillus subtilis* revealed the importance of membrane potential in response to translation inhibitors^26^. These new tools highlight the importance of membrane potential controlling bacterial physiology, and our ability to now study electrophysiology at the single cell level.

Despite the debate on the bactericidal mechanism of aminoglycoside, there is broad agreement that bacterial membrane potential plays a critical role. In this paper, we sought to investigate the influence of membrane potential in mediating bactericide upon treatment with aminoglycosides. We used live cell microscopy to maintain high spatial and temporal resolution while also resolving any heterogeneity within the population. We found that lethal concentrations of aminoglycosides induced voltage hyperpolarization leading to large fluctuations in cytoplasmic calcium that persisted for > 48 hours after treatment. We found these transients were correlated with the inability of cells to regrow, giving us a technique to measure the onset of cell death in real time at the single cell level. We found evidence that the transients arise from decreased ribosomal consumption of ATP leading to a reversal of the F1Fo-ATPase. The voltage hyperpolarization, in tandem with mistranslated proteins in the membrane, induced the bactericidal action. Our model proposes a new mechanism which links the chemical energy state of the cell with membrane potential dysregulation that can lead to death.

## Results

### Voltage is not necessary for aminoglycoside uptake or inner membrane pore formation in *E. coli* but is required for bactericidal activity

The proton ionophore cyanide m-chlorophenyl hydrazine (CCCP) dissipates voltage gradients, and is known to protect *E. coli* against the bactericidal activity and EDP-II uptake of aminoglycosides^5,6^. A colony forming unit (CFU) assay was performed using a glucose minimal medium (PMM, see methods and materials) in the presence of aminoglycosides. These measurements showed cells continued to grow in PMM in the presence or absence of CCCP (Fig 1A). Treatment of cells with aminoglycosides alone caused a rapid reduction in CFUs. In contrast aminoglycoside treatment of cells pre-treated with CCCP showed bacteriostatic activity (Fig 1A).

**Figure 1:**
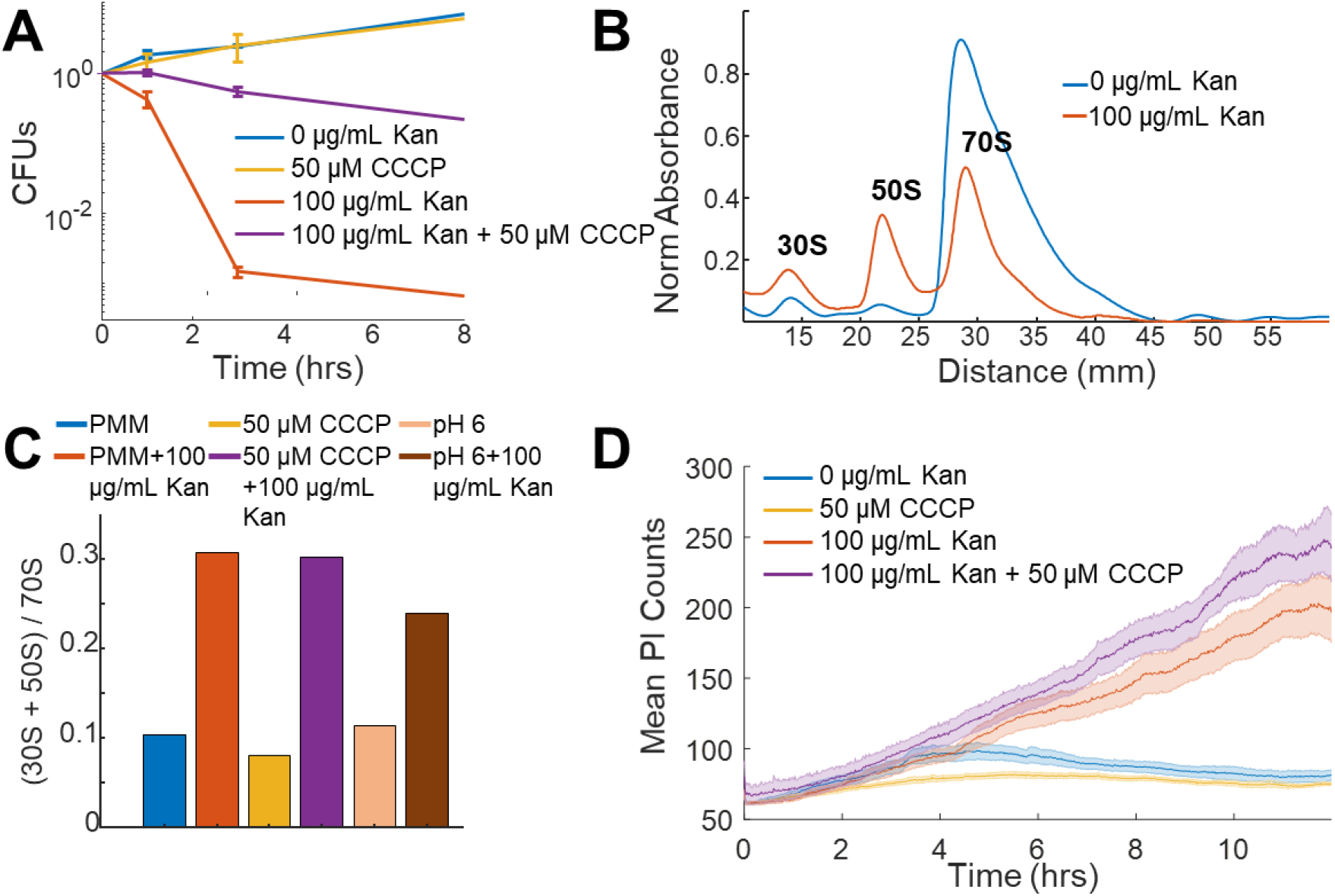
Voltage is not necessary for aminoglycoside uptake or inner membrane pore formation in *E. coli* but is required for bactericidal activity. (A) Colony forming units (CFUs) of untreated cells (blue) over four time points compared to cells treated with 50 μM CCCP (yellow), 100 μg/mL kanamycin (orange), and 50 μM CCCP + 100 μg/mL kanamycin (purple). Each curve averages 3 biological replicates, with mean and standard deviation plotted for each time point. (B) Ribosomal sucrose gradient depth plotted against 254 nm absorbance from LB grown *E. coli* treated with vehicle (blue), 100 μg/mL kanamycin (orange). The 30S, 50S, and 70S peaks are labelled. (C) Ratio of the area under the curve for the 30S + 50S to 70S peaks from *E. coli* in PMM pH 7.5, +50 μM CCCP, or pH 6 in the presence or absence of kanamycin. (D) Propidium iodide (3.75 μM in PMM) fluorescence in cells that were untreated (blue), 50 μM CCCP (yellow), 100 μg/mL kanamycin (orange), and 50 μM CCCP + 100 μg/mL kanamycin (purple) treated. The curve is the mean (solid) and standard deviation (shaded) for 3 biological replicates. Related to Supplementary Figure 1.

To more carefully examine the contrasting data that CCCP treated cells were growth inhibited in the presence of aminoglycoside, and the evidence that voltage is necessary for aminoglycoside uptake, a polysome analysis was used to assess ribosomal assembly in these conditions (Fig 1B)^27^. Untreated cells showed a majority of 70S particles, while addition of aminoglycosides caused a large fraction of ribosomes to split into 30S and 50S subunits^28^. Unexpectedly, ribosomes in aminoglycoside treated cells showed equal dissociation in the presence or absence of CCCP (Fig 1C, Fig S1A), despite the dramatic difference in drug activity. Aminoglycoside treatment at pH 6, which also has reduced membrane potential (see supplementary discussion), showed bacteriostatic activity and ribosomal dissociation (Fig 1C). In addition to chemical perturbations, naturally occurring mutations in bacterial populations can lead to protection against aminoglycosides arising from a decrease in membrane potential^2,17^. These mutations often occur in the electron transport chain, and reduce aminoglycoside uptake while concomitantly increasing survival^2^. Mutations of genes in the *nuo* operon have reduced uptake and death^2^, but have equivalent aminoglycoside induced ribosomal dissociation (Fig S1B). Though uptake of aminoglycosides in the absence of membrane potential has been observed^16^, the equivalent effect on ribosomal fraction abundance in *E. coli,* independent of voltage, had not been observed previously to our knowledge.

The clear uptake of aminoglycosides in the absence or alteration of membrane voltage suggested mistranslated proteins that induce membrane pores^13,14^ could also occur. We measured the uptake of propidium iodide (PI), a membrane impermeable DNA binding fluorescent dye, in the presence of aminoglycosides. The aminoglycoside treated population showed increasing PI fluorescence as compared to untreated cells (Fig 1D), indicating a loss of membrane integrity which correlated with the kinetics of cell death when measured by CFUs. Pre-treating cells with CCCP, however, showed a similar aminoglycoside induced increase in PI fluorescence, despite the switch from bactericidal to bacteriostatic activity. Chloramphenicol, a bacteriostatic translation inhibitor, induced only small increases in PI fluorescence (Fig S1C). These data suggested that protein mistranslation and membrane destabilization occur in the absence of membrane potential and are not sufficient to cause bactericidal activity. Given the discrepancy between CFUs, ribosomal dissociation, and PI uptake, we hypothesized voltage led to bactericide through mechanisms other than drug uptake. We therefore considered if bactericidal activity could arise through a combination of the mistranslated protein induced pore formation and membrane hyperpolarization. In order to test this hypothesis, we turned to single cell measurements of bacterial electrophysiology.

### Voltage and calcium exhibit altered electrophysiological flux in response to aminoglycosides

Fluorescent sensors of voltage and calcium have been used to monitor electrophysiology in bacteria at the single cell level with high time resolution^22,23,26,29^. We used the genetically encoded sensor, PROPS, to measure voltage dynamics after 2 hours of treatment with kanamycin. The aminoglycoside treated cells had larger fluorescent transients as compared to untreated cells (Fig S2A), but the high light intensities required prohibited long-term monitoring of single cells. GCaMP6, a fluorescent calcium indicator, is bright and sensitive enough to monitor live cells over hours or days, and we previously established calcium spikes were intrinsically linked to voltage fluctuations^23^. Individual *E. coli* expressing a fusion of GCaMP6f (calcium sensor) and mScarlet (spectrally independent control) were imaged upon exposure to 0 μg/mL or 100 μg/mL kanamycin and were monitored for 8 hours. Cells treated with antibiotic ceased growth and after ~2 hours showed large, non-oscillatory fluctuations which were uncoordinated between neighboring cells and not seen in untreated cells (Fig 2A, Supplementary Information (SI) Movie 1). Untreated *E. coli* had few cells with transients compared to drug treated cells and grew normally (Fig S2B,C, SI Movie 2). These drug induced transients were larger than previously observed mechanically induced fluctuations^23^. At a concentration of 30 μg/mL kanamycin >99.99% of cells cannot form colonies after six hours, yet we see transients >48 hours after kanamycin treatment at that concentration (Fig S2D,E). The delay between antibiotic exposure and the appearance of calcium transients varied across the population with a mean time of 1.64 hours after treatment (Fig S2F). The fraction of cells showing transients increased with increasing concentrations of kanamycin (Fig 2B). These data showed that aminoglycosides induced large electrophysiological effects that arise at similar timescales to cell death measured by CFUs.

**Figure 2:**
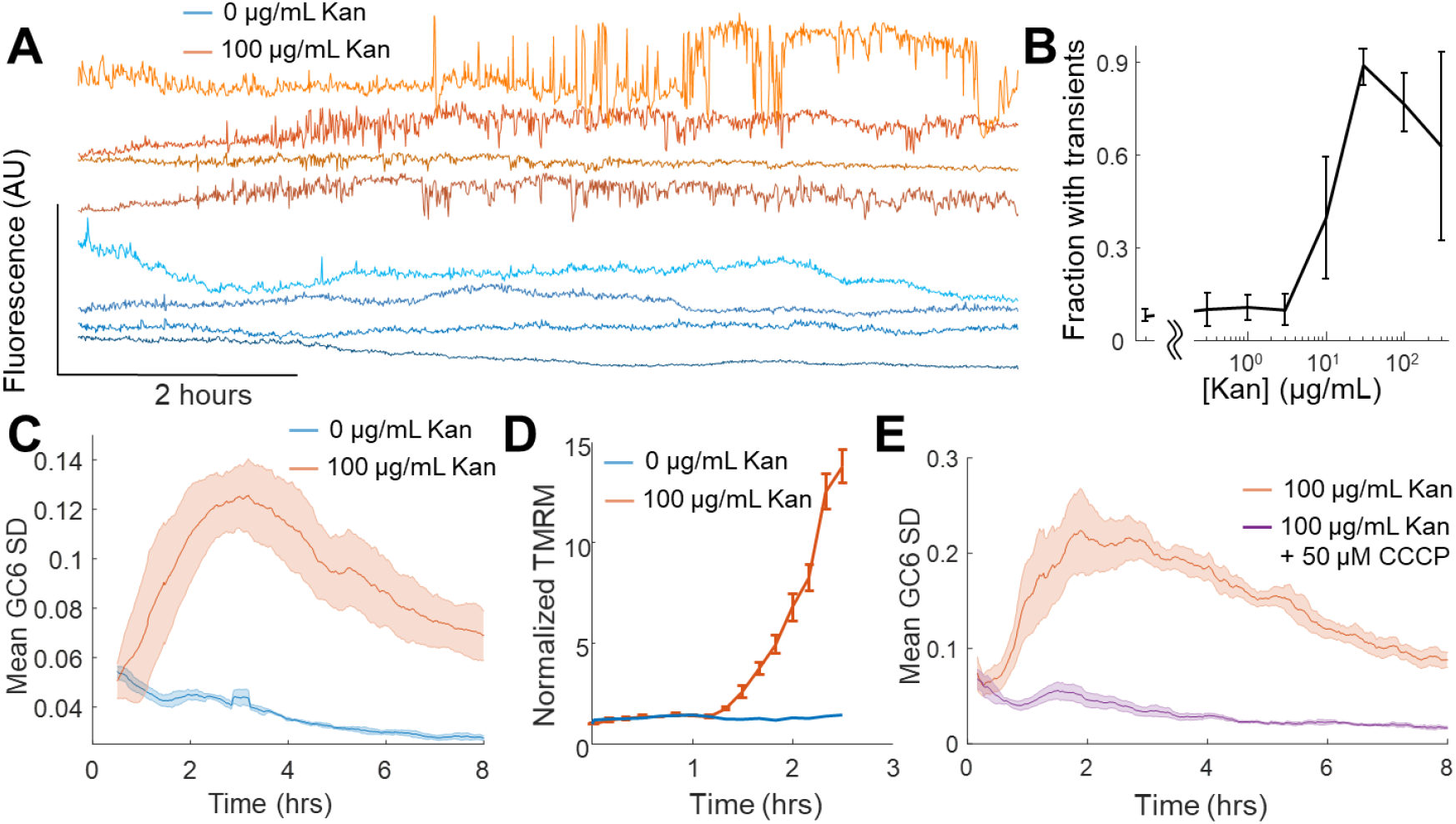
Voltage and calcium exhibit altered electrophysiological flux in response to aminoglycosides. Time traces of GCaMP6 fluorescence from single cells treated with 0 μg/mL (blue shades) and 100 μg/mL (orange shades) kanamycin. Individual cells display non-oscillatory transients. The fraction of cells in a population of GCaMP6F expressing cells *E. coli* experiencing the transients in A at different concentrations of Kanamycin. The mean (line) and standard deviation (error bars) are shown for three biological replicates. (C) The average (solid line) and standard deviation (shading) of the moving GCaMP6f standard deviation (SD) over time from 0 μg/mL (blue) and 100 μg/mL kanamycin (orange) treated cells. (D) TMRM fluorescence from untreated (blue) or kanamycin treated (orange) (100 μg/mL, 2 hours) *E. coli* measured by cytometry. The average (line) and standard deviation (error bars) of three biological replicates are plotted. (E) The average (solid line) and standard deviation (shading) of the moving GCaMP6f Standard Deviation over time from 100 μg/mL kanamycin treated cells in the absence (orange) or presence (purple) of 50 μM CCCP. Related to Supplementary Figures 2-5.

In order to compare the kinetics of the aminoglycoside response of populations of cells across treatment conditions, we needed a metric that would encompass the fluorescent dynamics across many cells. To visualize the transients across a population, a moving standard deviation was calculated for each cell, and then averaged across all cells. This mean of the moving standard deviation (taken from 30-500 cells) was considered one biological replicate, and the average and standard error of 3 biological replicates is then plotted (Fig 2C, Fig S3). This metric will depend strongly on the microscope system used, and thus requires relative comparisons of treated versus control under otherwise identical imaging conditions. We defined a drug induced calcium transient as any cell that showed a moving standard deviation (SD) >7-fold above untreated cells for >40 minutes. The GCaMP moving SD metric can separate treated and untreated populations of *E. coli*. All aminoglycosides tested exhibited a concentration dependent onset of calcium transients, as well as significantly increased GCaMP SD, but other bacteriostatic or bactericidal antibiotics had neither (Fig S4A,B). Our measurements do not rule out the possibility of other ions moving across the membrane^30^, and indeed we see that proton concentrations as measured by the red fluorescent pH indicator, pHuji^31^ also show transients, but their initial amplitude is much smaller than the calcium transients (Fig S4C,D). A lack of sufficient sensors prohibited us from measuring other ions at these temporal and spatial scales.

Given the observation that CCCP and low pH eliminated the calcium transients, we hypothesized that these large fluorescent changes were a product of a more polarized membrane potential, which would be consistent with the positive feedback of drug uptake model^2,8,14^. Tetramethylrhodamine methylester (TMRM), a membrane permeable fluorescent voltage reporter, accumulates in polarized mitochondria^32^ and *E. coli*^20,33^. Untreated *E. coli* showed no change in intracellular TMRM levels over 2.5 hours (Fig 2D). Cells treated with kanamycin showed a sharp increase in TMRM fluorescence after 80 minutes, corresponding to a change of −72 mV after 2.5 hours (see materials and methods). Assuming a resting potential of −150 mV the treated cells would have a membrane voltage of −222 mV. This observation is consistent with an aminoglycoside induced change in membrane potential occurring at the same time as the calcium transients.

If aberrant voltage induced the calcium transients, dissipating the voltage would eliminate the transients. Cells expressing GCaMP6 were treated with CCCP and compared to kanamycin exposure alone (Fig 2E). CCCP treated cells showed no increase in GCaMP6f SD, or individual calcium transients. Cells treated at pH 6 also showed no increase in calcium transients (Fig S5A) and showed no hyperpolarization measured by TMRM (Fig S5B). Knockouts of the *nuo* operon show altered kinetics in the onset of the GCaMP6f SD, as well as a lower amplitude in response to aminoglycoside treatment (Fig S4C). Together, these data show that aminoglycosides induced hyperpolarization and large ionic fluctuations only in the presence of membrane voltage, and that chemical or genetic alterations of membrane voltage affect the GCaMP6 response.

### Single cell calcium flux predicts cellular aminoglycoside response

The onset of voltage hyperpolarization, calcium transients, and cell death as measured by CFUs suggested the observed fluorescent calcium traces could be a good technique to measure bactericide at the single cell level. Fluorescence measurements were taken under continuous flow during the addition, then removal, of kanamycin. As expected, antibiotic exposure induced large calcium transients in many cells. After 4 hours of kanamycin exposure, medium without drug was added, and ~2% of cells reinitiated cell division (recovered cells, 35/1727 cells, Fig 3A, SI Movie 3). Of the 35 recovered cells, none exhibited drug induced calcium transients during or after antibiotic exposure (Fig 3B), and the population of recovered cells had lower calcium fluctuations as compared to arrested cells (Fig 3C). Recovered cells were not genetically resistant, as a second exposure to kanamycin stopped growth and induced calcium transients in daughter cells (Fig S6A-C). Finally, within an untreated population, a small fraction of cells exhibited transients (22 of 1544), where each cell with transients did not divide (Fig S6D-F). In all cases tested calcium transients correlated with reduced population viability; conditions with fewer calcium transients increased CFUs, and any cell that exhibited transients did not regrow. This data provided a technique to measure one hallmark of single cell death in *E. coli* in real time as all observations of these transients indicated that a cell experiencing them was rendered unable to divide, though we are not able to definitively say that the transients caused cell death.

**Figure 3:**
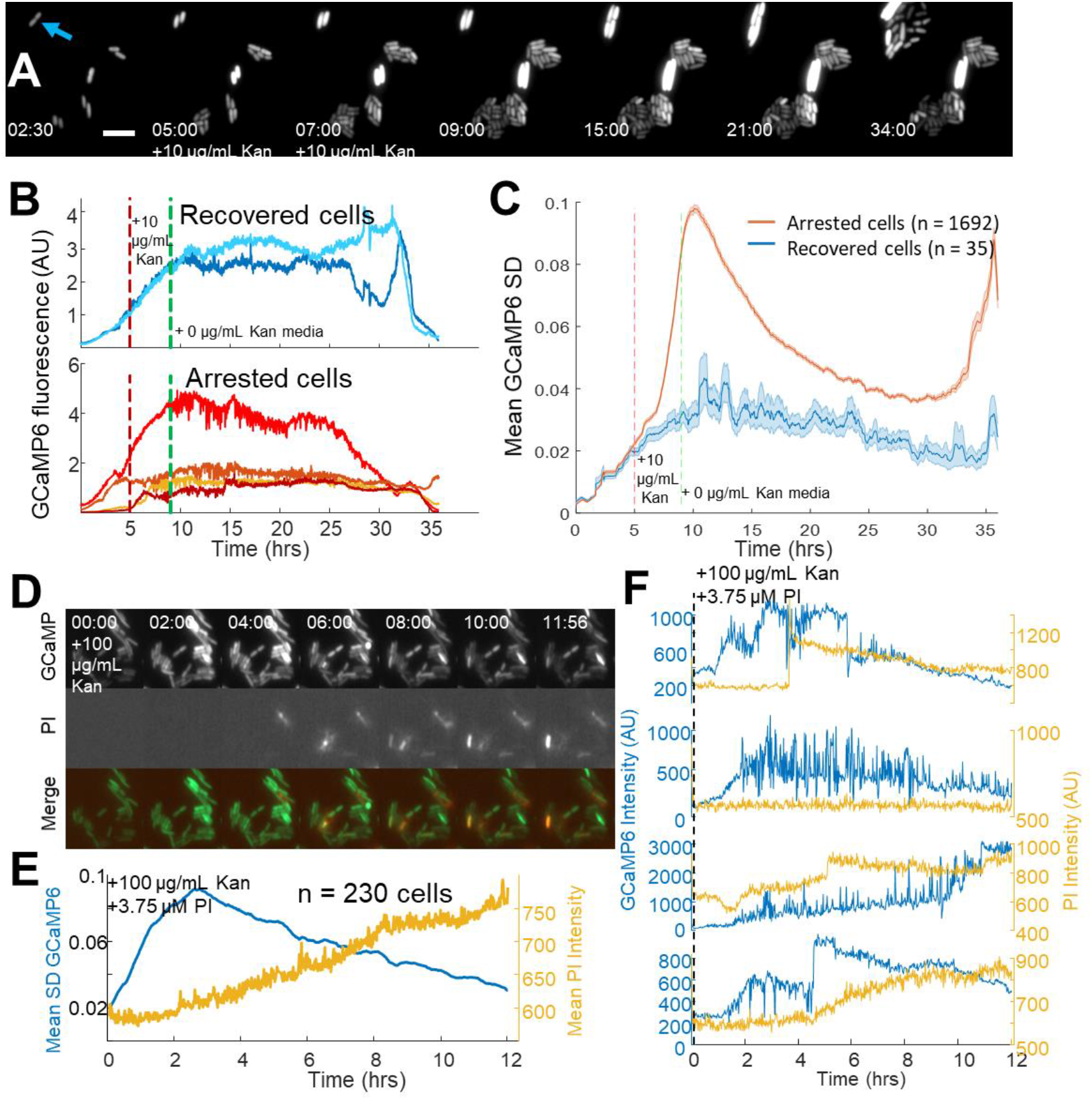
Single cell calcium flux predicts cellular aminoglycoside response. (A) Strip chart of cells expressing GCaMP6f. Cells were imaged in PMM alone for 5 hours, then exposed to 10 μg/mL kanamycin for 4 hours. After 4 hours, PMM alone was flowed in for an additional 26 hours. The blue arrow indicates a cell that was able to divide after treatment with kanamycin, 5 μm scale bar. Time is shown in (HH:MM) format. (B) Individual GCaMP6 time traces from cells that regrow after treatment compared to a random selection of cells that do not regrow within 24 hours. (C) The average (line) and standard deviation (shaded region) of the moving SD from all cells that regrow (blue) vs those that do not regrow (red). (D) Strip chart showing GCaMP6 fluorescence (top), propidium iodide fluorescence (middle), and the merge. Cells were treated with 100 μg/mL kanamycin at time t = 0. Time is shown in (HH:MM) format. (E) The mean GCaMP6 standard deviation for the population is shown in blue. Yellow shows the population average of the PI fluorescence. (F) Time traces of individual cells showing the GCaMP6 fluorescence (blue) and the PI fluorescence (yellow) on the same cells. The PI fluorescence was not correlated with the onset of transients, though many cells did uptake PI during the course of the experiment. Related to Supplementary Figure 6.

Spectrally separating PI and GCaMP enabled us to study the kinetics between catastrophic calcium transients and pore formation in single cells. The mistranslation that causes pore formation was previously measured to occur within a half hour of aminoglycoside treatment^14^. We hypothesized that mistranslated proteins in the plasma membrane created an ionic imbalance in polarized cells leading to the observed calcium transients. To test our hypothesis, we incubated GCaMP6 expressing *E. coli* with PI in the presence of aminoglycoside (Fig 3D). The population average showed a smoothly increasing level of PI uptake upon aminoglycoside exposure (Fig 3E), similar to our previous data. However, the GCaMP6 moving SD increased well before appreciable PI uptake. Individual cells showed calcium transients preceded PI entry into the cytoplasm, and that PI often increased in very large bursts (Fig 3F). Thus, pores large enough to accommodate PI occurred after aminoglycoside induced hyperpolarization and catastrophic calcium transients, suggesting bactericidal activity occurred prior to pore formation.

### Voltage toggles between bactericidal and bacteriostatic activity in aminoglycoside treated cells

The data above showed that aminoglycoside uptake, ribosome dissociation, and mistranslated protein can occur without membrane potential. Aminoglycosides in the absence of a voltage exhibited a bacteriostatic effect, but voltage induced bactericide. We therefore sought to explore the requirements of voltage as the bactericidal keystone in *E. coli* by using the calcium transients as a real time marker of permanent cell cycle arrest, while controlling the chemical environment to actuate membrane voltage.

Treating cells with aminoglycoside induced calcium transients (Fig 4A top, Fig S7A top) as expected. However, removing the voltage either through addition of CCCP or lowering pH immediately ceased all transients at the single cell and population levels (Fig 4A,B, Fig S7A,B), though no cells re-initiated cell division. Thus, voltage was necessary for the calcium transients to occur. Conversely, *E. coli* was incubated with kanamycin in the presence or absence of CCCP for 4 hours and showed calcium transients only in the cells without CCCP as expected (Fig 4C top). Removal of kanamycin and CCCP initiated transients within 7 minutes, much faster than the appearance of transients from aminoglycoside treatment without CCCP (Fig 4C, D). Similar results were seen exchanging pH 6 with pH 7.5 to reestablish a membrane voltage (Fig S7C,D). The rapid onset showed that aminoglycosides can exert bactericidal activity immediately upon reestablishment of membrane voltage, and that in the conditions tested, voltage is sufficient to induce catastrophic calcium transients which were correlated with cell death.

**Figure 4:**
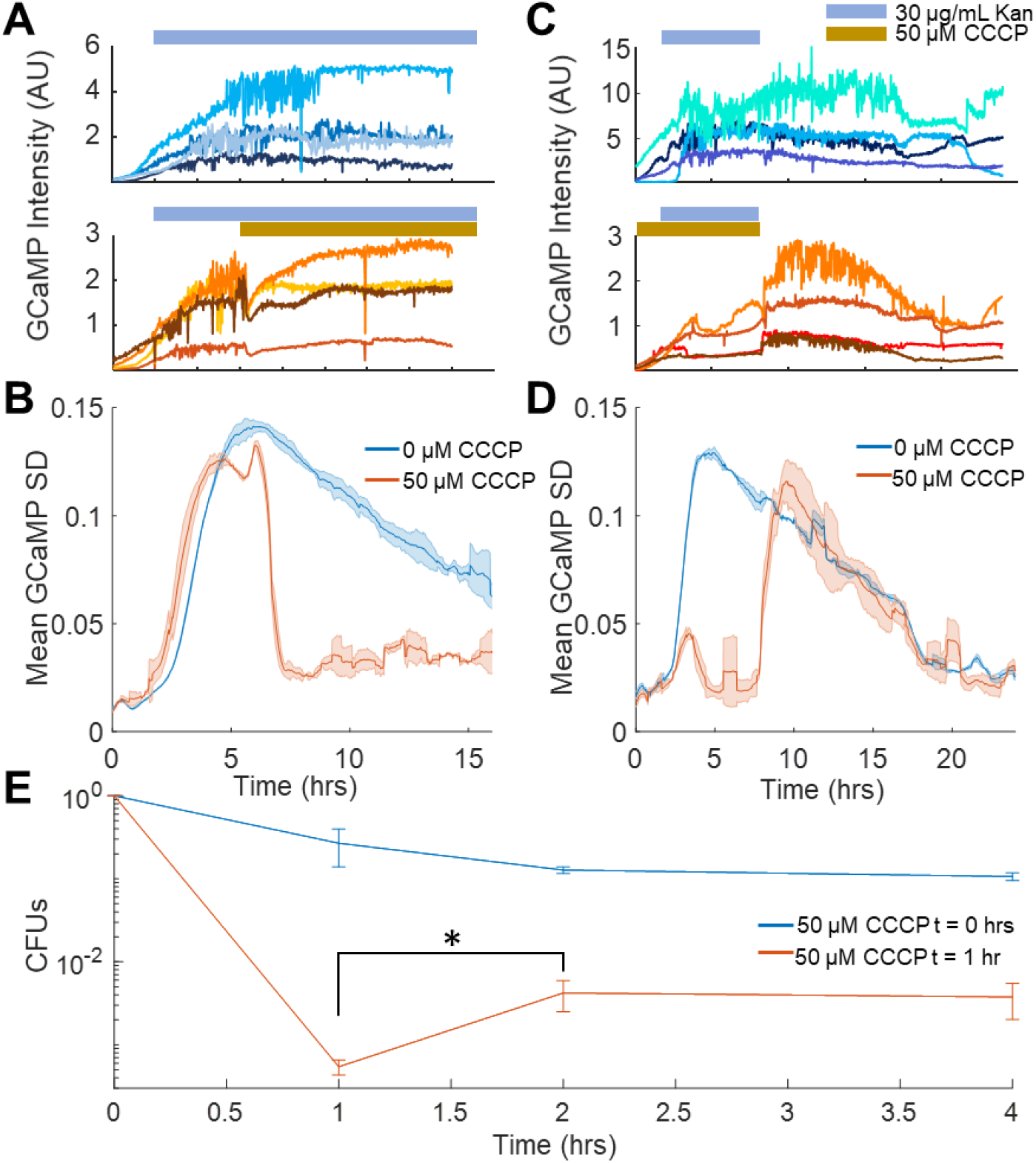
Voltage toggles between bactericidal and bacteriostatic activity in aminoglycoside treated cells. (A) Single cell traces of GCaMP6f intensity over time upon treatment with kanamycin (orange bar, top), or with kanamycin followed by CCCP (yellow bar, bottom). (B) Mean GCaMP6f moving SD of biological replicates over time. The population traces are the mean of the single cell experiments in B, with kanamycin (blue, corresponds to A top) or kanamycin +CCCP treated cells (orange, corresponds to A bottom). (C) Single cell traces of GCaMP6f intensity over time after kanamycin was flowed across the cells at 2 hours that were pretreated with vehicle (top) or CCCP (bottom). Kanamycin and CCCP were then flowed out of the chamber at 6 hours. (D) Mean GCaMP6f moving SD of biological replicates over time. The population traces are the mean of the single cell experiments in C, with kanamycin (blue, corresponds to C top) or kanamycin +CCCP treated cells (orange, corresponds to C bottom). (E) Cells treated with 10 μg/mL gentamycin with CCCP added at t = 0 hrs (blue line), or CCCP added at t = 1 hr (orange line). CFUs taken at 1 hour were counted before addition of CCCP, which increased the number of surviving cells at t = 2 hrs. *p < 0.05. Related to Supplementary Figure 7.

If voltage hyperpolarization induced cell death, a prediction is that chemically removing voltage before the onset of transients would protect cells, even if the cells are maintained in the presence of aminoglycoside. If cells were treated with aminoglycoside, followed by CCCP addition, there would be an increase in the number of surviving cells, even if those cells were maintained in the antibiotic for a longer period of time. To test this prediction, *E. coli* were treated with 10 μg/mL gentamicin and CFUs were counted at 60 minutes. At that time, CCCP was added to the medium, and cells were incubated for another 60 minutes with aminoglycoside and CCCP. After 2 hours, CFUs were counted again, and there was a 22x increase in CFUs as compared to the 1-hour time point (Fig 4E). This data shows that the conditions for cell death had been established at 1 hour and that cells then plated onto LB would still die. However, cells treated with CCCP at 1 hour avoided the hyperpolarization induced calcium transients and had a correspondingly higher survival rate.

### ATP dysregulation precedes voltage induced bactericidal killing

Published evidence suggests that metabolic dysfunction correlates with translation inhibitor efficacy^34–36^. This was hypothesized to be associated with bacterial energetic investment in protein production^37^. Furthermore, a reduction in ribosome concentration has been annotated as a means to protect persister cells^38^. We reasoned that the sudden change in energetic demand from the loss of a large fraction of 70S translating ribosomes could free up ATP and GTP to be used in other processes. To connect this shift in energetics to aminoglycoside induced voltage dysregulation, we considered how *E. coli* generate a membrane voltage in aerobic environments. In the presence of glucose, *E. coli* use glycolysis to power the NADH dehydrogenase assembly (Complex I) and induce a proton motive force (PMF). The F1Fo-ATPase then depletes the PMF to generate ATP. However, the F1Fo-ATPase can be run in reverse, using ATP hydrolysis to generate a membrane voltage, which occurs in anaerobic conditions to power flagellar rotation^39^. We hypothesized that aminoglycosides increased cellular ATP flux through non-ribosomal sinks, leading to hyperpolarization via the combined activity of the NADH dehydrogenase and a reversed F1Fo-ATPase.

We initially measured ATP concentration in *E. coli* using a ratiometric fluorescent ATP sensor, mRuby-iATPSnFR1.0^40^. Gentamicin treatment increased the 488/561 nm fluorescence ratio by 50% within 2 hours of treatment (Fig 5A). Cells at low pH or in the presence of CCCP also showed ATP increases expected from ribosome dissociation (Fig S8A,B). Other non-aminoglycoside translation inhibitors which exhibit bacteriostatic activity also showed increasing ATP (Fig S8C). Consistent with our observation that recovered cells did not exhibit calcium transients, cells that recovered after 4 hours of kanamycin treatment had lower ATP compared to arrested cells (Fig 5B). We attempted to quantify the absolute change in ATP concentration in populations of cells, as our single cell data indicated that ATP levels were increased when cells were treated with aminoglycosides. Using a luminescence-based assay, we determined that steady state levels of ATP in gentamicin treated *E. coli* were significantly lower than untreated controls (Fig S8D) in the first half hour of treatment, which was inconsistent with our iATPSnFR single cell data. This data is, however, consistent with an increased ATP flux through consumers other than ribosomes, such as the F1Fo-ATPase. We suspected that the genetically encoded ATP sensor can act as a buffer absorbing some of this ATP flux from a loss of translation, while the luminescence-based assay measures absolute values after the cells are permeabilized. This interpretation is consistent with recent results which show an increase in an alarmone with an ATP precursor after aminoglycoside treatment^41^. Collectively these data suggest that there may be a change in metabolic flux in the system, and are consistent with prior observations of aminoglycoside treated cells, which were found to leak NTPs^6^ and increase respiration^18^. This change in ATP flux is consistent with a number of other observations in the field correlating metabolism with translation inhibitor efficacy^34–36,42^.

**Figure 5:**
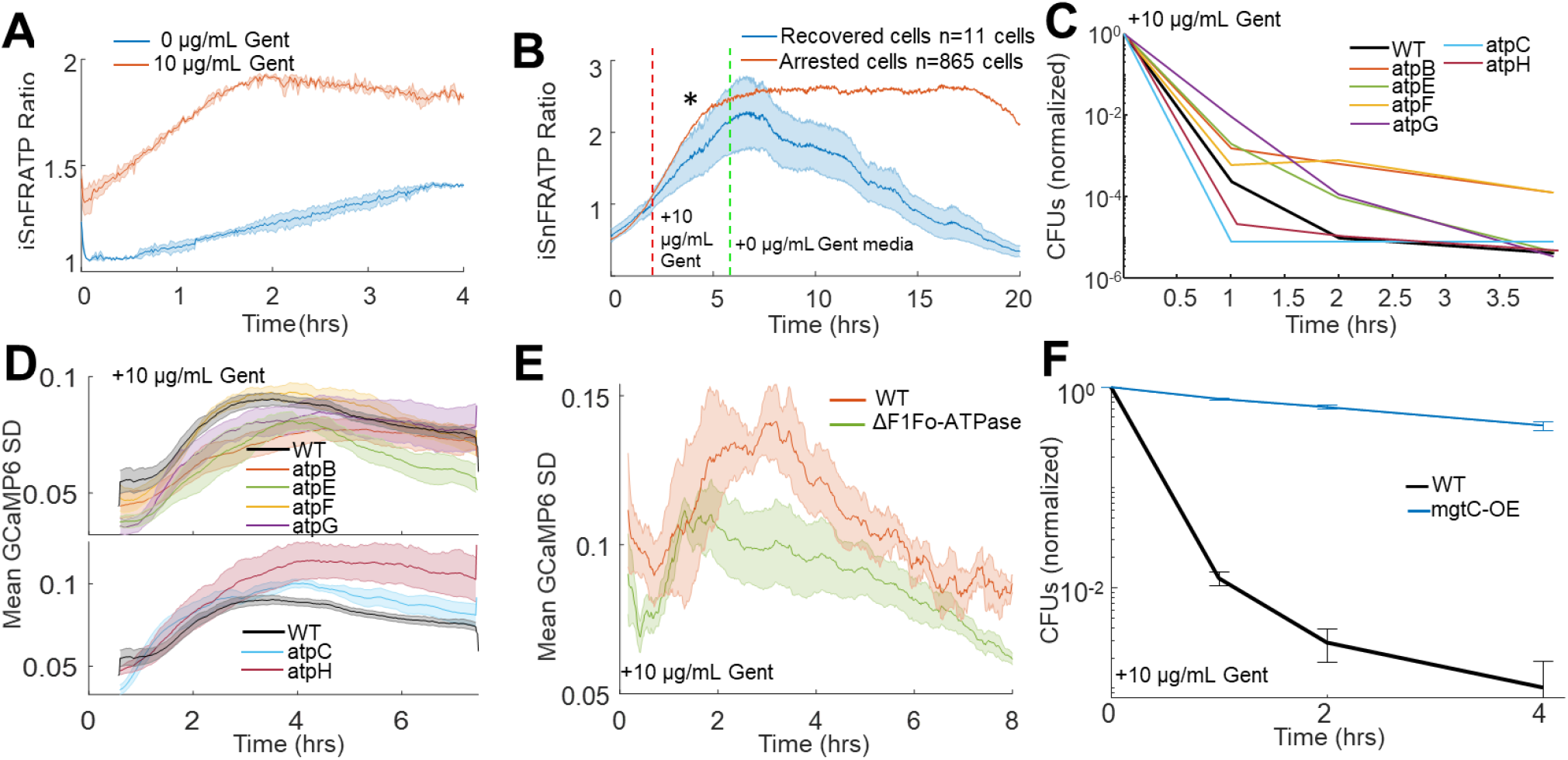
ATP dysregulation precedes voltage induced bactericidal killing. (A) iATPSnFR ratios from *E. coli* treated with vehicle (blue) or 10 μg/mL gentamicin (orange). The ratio of iATPSnFR (488 nm) to mRuby (561 nm) indicates ATP concentration. Each trace averages 2 biological replicates. (B-F) Cells treated with 10 μg/mL gentamicin. (B) iATPSnFR ratios from gentamicin treated cells that do (blue) or do not (orange) regrow. The star represents a significance of < 0.05 tested at 2 hours after treatment using a student t-test with unequal variance. (C) Normalized CFUs of gentamicin treated knockouts of components of the F1Fo-ATPase compared to WT. Each data point is in biological triplicate. (D) Mean moving GCaMP6f SD for gentamicin treated F1Fo-ATPase component knockouts compared to WT. Each curve averages 4 biological replicates. (E) Mean moving GCaMP6f SD for gentamicin treated *E. coli* strain DK8, missing all components of the F1Fo-ATPase compared to WT. Each curve averages 4 biological replicates. (F) Normalized CFUs of gentamicin treated mgtC expressing *E. coli* compared to WT. Related to Supplementary Figure 8.

If aminoglycosides induced ATP hydrolysis and hyperpolarization via the F1Fo-ATPase, then pump component knockouts should reduce calcium transients, show increased CFUs compared to WT, yet also show increased ATP due to the absence of hydrolysis. Knockouts from the proton conducting Fo domain (atpB, atpE, atpF) as well as atpG had increased CFUs and reduced calcium transients compared to WT (Fig 5C,D top), and all tested ATPase knockouts showed gentamicin induced ATP accumulation (Fig S8E). Interestingly knockouts of atpC (ε-subunit), which has increased gentamicin sensitivity^43^, and atpH (δ-subunit) both decreased the time to calcium transient onset and reduced CFUs faster than WT (Fig 5C,D bottom). AtpC biases the motor in the direction of ATP production^44^, while atpH acts as a filter for proton conduction through the Fo domain^45^, thus the knockouts of these proteins would improve proton conduction through the F1Fo-ATPase thereby increasing membrane potential that can be generated by this pump, which are consistent with knockouts showing more rapid cell death. Furthermore, gentamicin treated Fo domain knockouts reduced hyperpolarization while, as their function predicts, atpC and atpH increased hyperpolarization relative to WT (Fig S8F).

Completely eliminating the F1Fo-ATPase^46^ also showed reduced calcium transients as compared to a strain with intact F1Fo-ATPase activity (Fig 5E). Finally, expression of a virulence factor from *Salmonella,* mgtC, eliminated the bactericidal activity of aminoglycosides in *E. coli* (Fig 5F). MgtC is an inhibitor of the F1Fo-ATPase^47^, and aids in *Salmonella* infection and survival at low magnesium^48,49^. These data were all consistent with aminoglycosides inducing membrane hyperpolarization from ATP hydrolysis via the F1Fo-ATPase, ultimately leading to cell death.

## Discussion

Aminoglycosides are well established to bind and exert pleiotropic effects on ribosomes^5,50,51^, and numerous reports highlighted the importance of maintaining a membrane potential in aminoglycoside activity. This evidence included voltage dependent aminoglycoside uptake^10^ and cell death correlated with the citric acid cycle and carbon source^35,52,53^. Metabolic changes can likewise induce changes in membrane voltage and the overall proton motive force. The relationship between metabolism, proton motive force, and membrane potential has been typically seen as being requisite to the *uptake* of aminoglycosides, which was synonymous with cell death^5,11^. Our work has shown that membrane voltage is not essential for drug uptake, but rather the voltage is required to initiate the bactericidal mechanism after ribosome dissociation. Though we show a correlation between the ionic imbalance (calcium and pH transients) and cell death, we did not definitively prove they cause cell death, but rather they provide a convenient metric for cell death at the single cell level. Our data also does not preclude a mechanism of voltage enhanced aminoglycoside uptake^2,5^. Rather our work suggests that the uptake of aminoglycosides in the absence of a membrane potential is sufficient to create intracellular conditions, including ribosome dissociation, metabolic dysfunction, and pore formation, that allow the presence of a membrane potential to exert bactericidal effects. Our data is also consistent with other translation inhibitors hyperpolarizing membrane potential correlated with subsequent cell death^26^. We provide evidence that one mechanism by which this hyperpolarization can occur is through F1Fo-ATPase activity. We observed enhanced aminoglycoside killing in the strain atpC::kanR, which is missing the F1Fo-ATPase ε-subunit that typically biases the rotor in the direction of ATP synthesis. This observation suggests that F1Fo-ATPases with a higher likelihood of ATP hydrolysis enhance aminoglycoside killing, which would stem from the already ribosome-related dysregulation of metabolism. We observed similar enhanced aminoglycoside killing in the strain atpH::kanR that encodes the δ-subunit of the F1Fo-ATPase, which is able to block proton conduction^45^ and ATP hydrolysis^54^. Together, these data suggest that the difference between bactericidal activity of aminoglycosides compared to the bacteriostatic activity of other translation inhibitors may be the lack of the mistranslated membrane proteins causing pore formation. We hypothesize this mechanism kills bacteria by eliminating ion homeostasis in the presence of a membrane potential and pores that can leak ions. However, we currently lack tools to be able to induce the calcium transients in the absence of aminoglycosides, though perhaps channel rhodopsins will be able to mimic these effects.

One fascinating facet that remains to be explored is the period after aminoglycoside treatment that cells cannot divide but remain metabolically active for at least 2 days. If these arrested cells can still export quorum sensing molecules, they could send paracrine signals to untreated cells, and influence their behavior. This observation became clear by using sensitive genetically encoded fluorescent proteins, and these tools open up a new avenue to study the long-term effects of antibiotic treatment on cells and mixed cultures. Another curious corollary is the observation that protonophores enhance aminoglycoside killing in *Pseudomonas* biofilms^55,56^, which stands in opposition to our observation that protonophores protect planktonic *E. coli*. The differences driven by these species specific and context dependent observations will hopefully add to a more complete picture of aminoglycoside activity in multiple bacterial species.

The model of aminoglycoside induced death proposed from this work is consistent with evidence from other groups previous work, requires the presence of membrane pores and membrane potential to drive aminoglycoside bactericidal activity. Aminoglycosides enter the cell through an unknown mechanism, possibly through channels such as mscL^57^, which occurs long before a loss of membrane integrity. Once aminoglycosides enter the cell they bind ribosomes, disrupt a majority of translating 70S particles and cause mistranslation of protein^13,30^. As soon as ribosome disruption occurs, respiration^18^ and metabolism^34^ go through a substantial shift in flux. This disruption of metabolism enables non-canonical generators of membrane potential, such as the F1Fo-ATPase to drive changes in membrane potential. Why voltage is so toxic in the presence of the mistranslated membrane proteins remains to be explored, however this shift in understanding the role voltage plays in aminoglycoside lethality will hopefully provide a necessary rethinking of how these antibiotics function so much more effectively than other translation inhibitors. The difference between these mechanisms of bactericide and stasis could lead to novel antibiotics that impinge on the aminoglycoside mechanism of action.

## Supporting information

Supplemental figures discussion movie legends

## Acknowledgments

Thanks to Annette Erbse and Keda Zhou for help with ribosomal profiling, Theresa Nahreini for help with cytometery, and Karolin Luger, Joe Falke, Tom Cech and the Biochemistry Shared Instruments Pool at the University of Colorado Boulder for equipment resources. We thank Stephanie Bueler and John Rubinstein for the DK8 strain and helpful discussion. Thanks to Toshiharu Suzuki for the pTR-ASDS plasmid. We had helpful discussions with Corrie Detweiler, Amy Palmer, Ben Dodd, Stacey Scott, and Rose Luder. Special thanks to Thomas Yao for many helpful discussions.

## Funding

Searle Scholars Program and NIH New Innovator (1DP2GM123458) to J.M.K., T32 training grant (T32GM065103) and HHMI Gilliam Fellowship for Advanced Study to G.N.B. Flow cytometer was acquired with an instrumentation grant (NIH S10OD021601).

## Notes

### Competing Interest Statement

The authors have declared no competing interest.

### Summary of Updates

Figures now include drug and concentration when added. Supp Movie 3 now indicates the addition of kanamycin.

