## Supplemental figures discussion movie legends for "Membrane voltage dysregulation driven by metabolic dysfunction underlies bactericidal activity of aminoglycosides"

### Supplementary Information

**Supplementary Materials:**

Supplementary Discussion

Supplementary Figures 1-8

Supplementary Movies 1-3

Resources Table

Materials and methods

**Supplementary discussion**

*E. coli* energize their membrane through a proton motive force (PMF) that powers their flagellar motors and several membrane pumps. The PMF is the amount of free energy gained by a proton moving from one side of the membrane to the other, and the energy can be gained either by changes in pH (proton gradient) or voltage (membrane potential). The Nernst equation sets an equivalence between changes in pH and voltage as:

$$PMF = \Delta\psi * -58mV(\Delta pH)$$

*E. coli* typically try to maintain a cytoplasmic pH around 7.5, so that a changing extracellular pH will induce a corresponding change in the PMF. For example, if the extracellular pH is at pH 7.5, then there is no pH difference, so all of the PMF will be carried in the voltage component, which would be accomplished by establishing ionic gradients using pumps and channels. On the other hand, if the extracellular pH is low, for example pH 6, then the PMF could have a value -87 mV without having to maintain any voltage component. The PMF could be carried entirely by the change in pH, which could drive the flagellar motors and other PMF dependent processes in the membrane.

Thus, by changing the environmental pH from 7.5 to 6, we can lower the membrane voltage by fact that the cell will utilize the pH component of PMF without the need to generate an external voltage from other ions.

#### Supplementary Figures

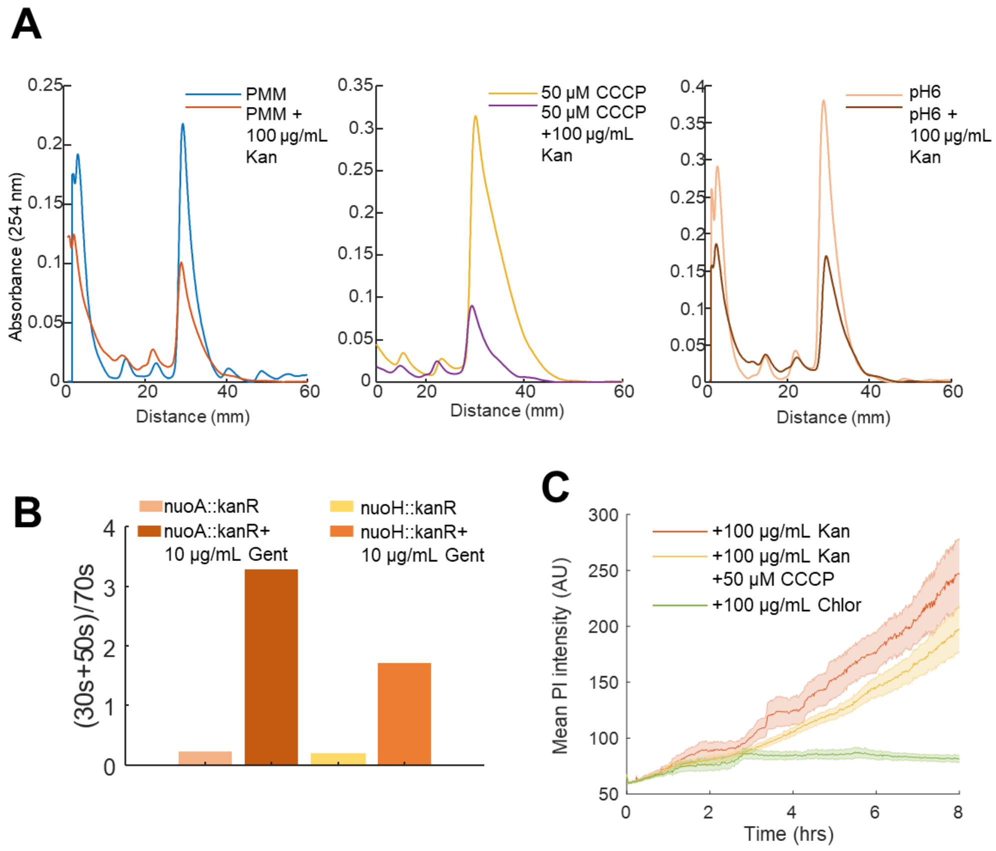


**Supplementary Figure 1. Related to Figure 1.**

(A) Ribosomal sucrose gradient depth plotted against 254 nm absorbance from *E. coli* in treatment conditions from Fig 1C. (B) Ratio of the area under the curve for the 30S + 50S to 70S peaks from nuoA::kanR and nuoH::kanR *E. coli* strains in the absence and presence of gentamicin. (C) The uptake of 3.75 µM propidium iodide (PI) was measured by microscopy in cells that were 100 µg/mL kanamycin (orange), 100 µg/mL kanamycin + 50 µM CCCP (yellow), and 100 µg/mL chloramphenicol (green) treated. The mean (line) and standard deviation (shaded) are plotted over time.

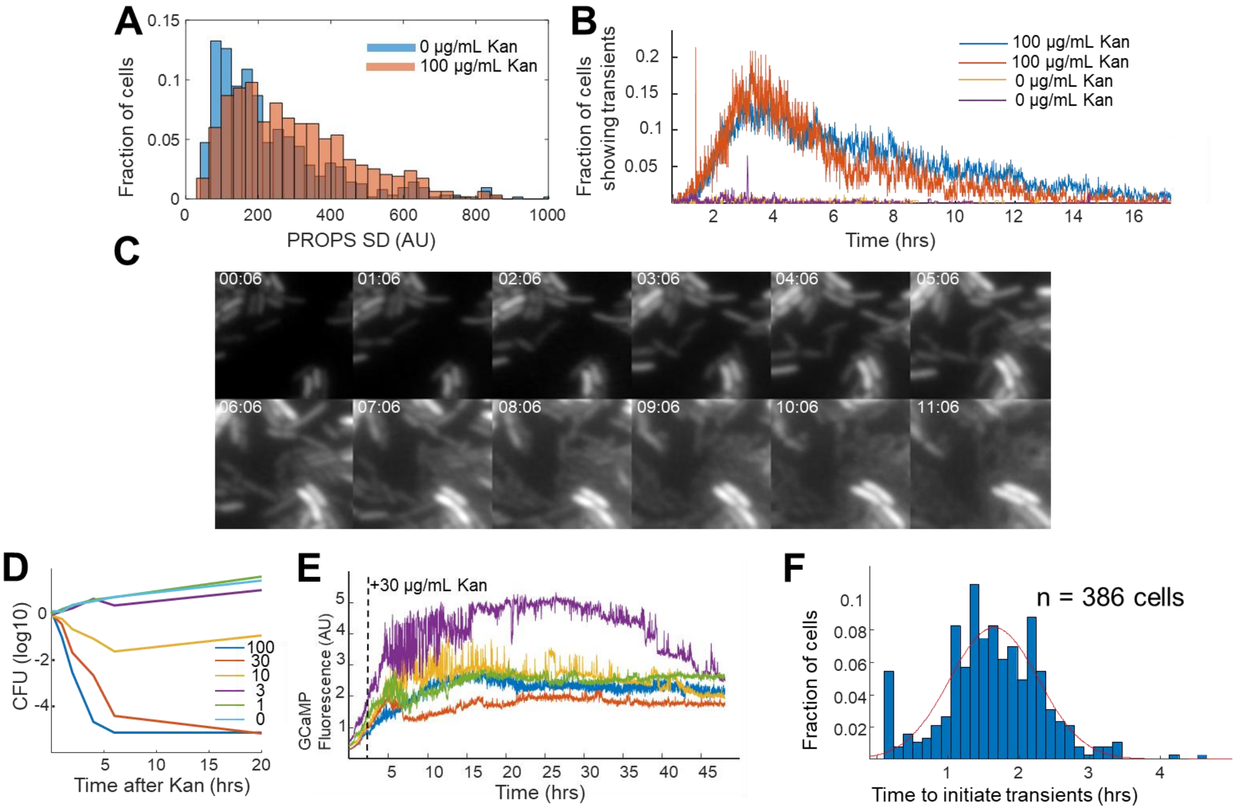


**Supplementary Figure 2. Related to Figure 2.**

(A) A histogram of the fraction of cells with a given standard deviation of PROPS fluorescence in the absence (blue) and presence (orange) of 100 µg/mL kanamycin. (B,C) Untreated cells have substantially fewer large calcium transients. (B) Fraction of cells exhibiting calcium transients as a function of time for untreated (yellow, purple) compared to treated (red, blue) cells. Each trace represents the mean of biological replicates of > 200 individual cells. Untreated cells do not show any coordinated transients. (C) Strip chart showing the growth of untreated cells. The time is shown in (HH:MM) format. (D) Survival of cells in liquid culture upon increasing kanamycin concentration in µg/mL, measured by CFUs at indicated dose and time. Each curve averages 3 biological replicates. (E) Kanamycin treated cells are capable of exhibiting transients for at least 48 hours. Time traces of GCaMP6 fluorescence from individual cells. Each color represents the mean fluorescence from a single cell. The black dotted line represents the addition of 30 µg/mL kanamycin. The majority of calcium transients occur within the first 12 hours after treatment, but individual cells still show activity even up to 48 hours after treatment. (F) Histogram shows the time of the first transient for cells that had transients. The population fit to a Gaussian with a peak of 1.64 hours after kanamycin addition.

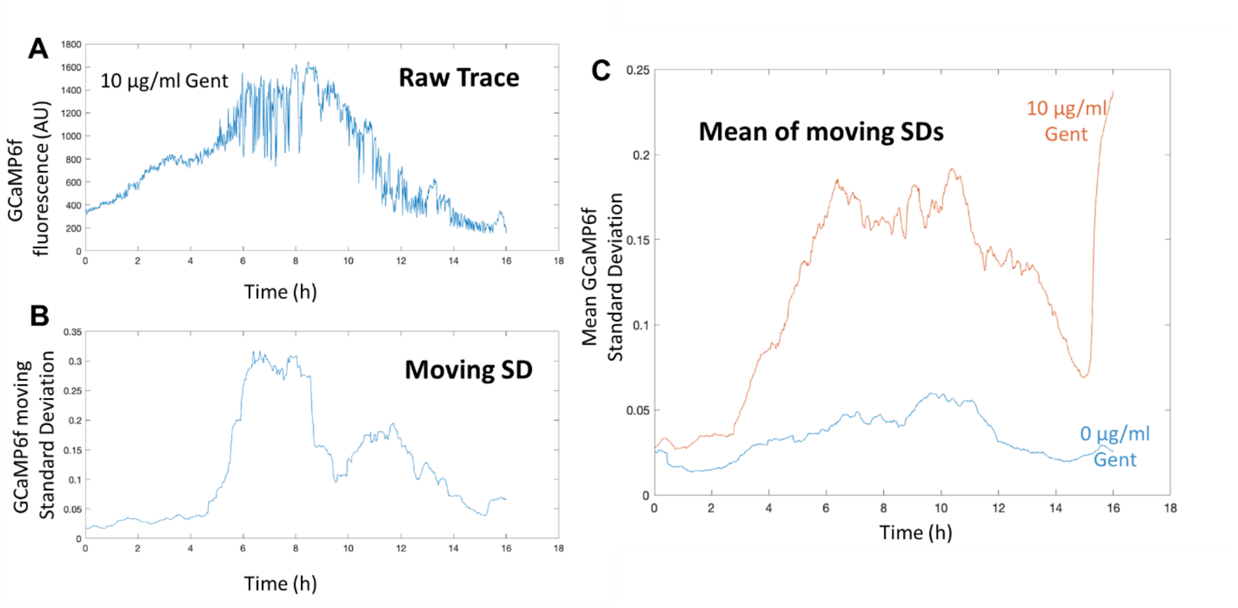


**Supplementary Figure 3. Related to Figure 2.**

(A) A time trace of GCaMP6f fluorescence from a single *E. coli* cell that has been exposed to 10 µg/mL of gentamicin. (B) The fluorescence changes over time in A were transformed to the GCaMP6f moving standard deviation by first normalizing by dividing by a 45-minute moving median. Then the moving standard deviation was calculated from the normalized trace using a 30-minute sliding window, the output of which is shown here. Large changes in the fluorescence changes in A (6 hours post treatment spiking) reveal themselves as a large change in B (a ~10-fold increase in GCaMP6f moving SD at the corresponding 6-hour mark compared to the first 4 hours post treatment). (C) The average of ~XX cells moving GCaMP6f moving standard deviation for 0 and 10 µg/mL gentamicin treatment reveal that this metric can reliably distinguish populations of treated and untreated cells.

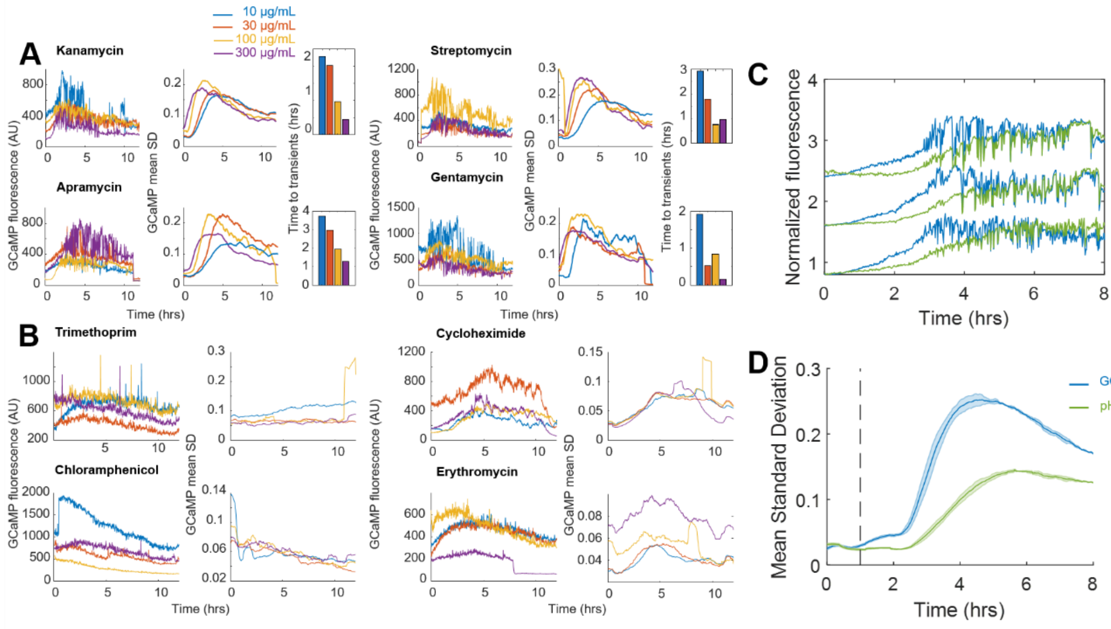


**Supplementary Figure 4. Related to Figure 2.**
(A) All aminoglycosides tested (kanamycin, streptomycin, apramycin, gentamicin) exhibit calcium transients in a concentration dependent fashion. For each compound tested, the fraction of cells exhibiting transients increased, while the time of transient onset decreased with increasing concentration of aminoglycoside treatment. Each three panel set of the compounds; Kanamycin, Streptomycin, Apramycin, and Gentamicin: (left) representative GCaMP6 time traces at 100 µg/mL compound, (middle) the mean GCaMP6 standard deviation of the population, and (right) the mean of the Gaussian fit for the onset of transients for a population as a function of treatment concentration. (B) Antibiotics that are not aminoglycosides did not induce catastrophic calcium transients in any fashion. For trimethoprim, cyclohexamide, chloramphenicol, and erythromycin, representative traces of individual cells are shown at a treatment of 100 µg/mL (Left). The mean GCaMP6 standard deviation for the population is shown across a titration of each antibiotic (right). Each trace is the mean of 2 biological replicates. (C) Single cell traces over time of pHuji (green) + GCaMP6f (blue) expressing cells treated with 30 µg/mL kanamycin. (D) Mean moving standard deviation plots of populations of cells represented in C. Dashed line represents addition of aminoglycoside treatment.

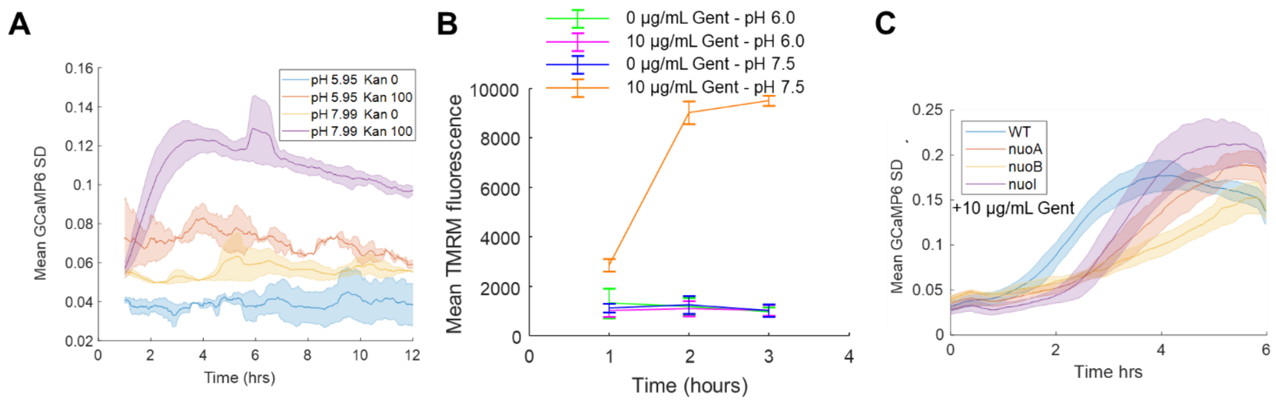


**Supplementary Figure 5. Related to Figure 2.**

(A) The average of the moving SD from a population of cells at pH 5.95 and pH 7.99, either treated with 0 or 100 µg/mL kanamycin. Each curve averages 3 biological replicates. Shading around the line is the standard deviation. (B) Mean (line) and standard deviation (error bars) of TMRM fluorescence of cells over time measured by cytometry in 200 nM TMRM in PMM. *E. coli* in pH 6.0 in the absence (green) or presence (magenta) of 10 µg/mL gentamicin were compared to *E. coli* at pH 7.5 in the absence (blue), and presence (orange) of 10 µg/mL gentamicin. (C) The average of the moving SD from a population of cells treated with 10 µg/mL gentamicin that were WT (blue), nuoA::kanR (orange), nuoB::kanR (yellow), and nuoI::kanR (purple) strains. Each curve averages 3 biological replicates. Shading around the line is the standard deviation.

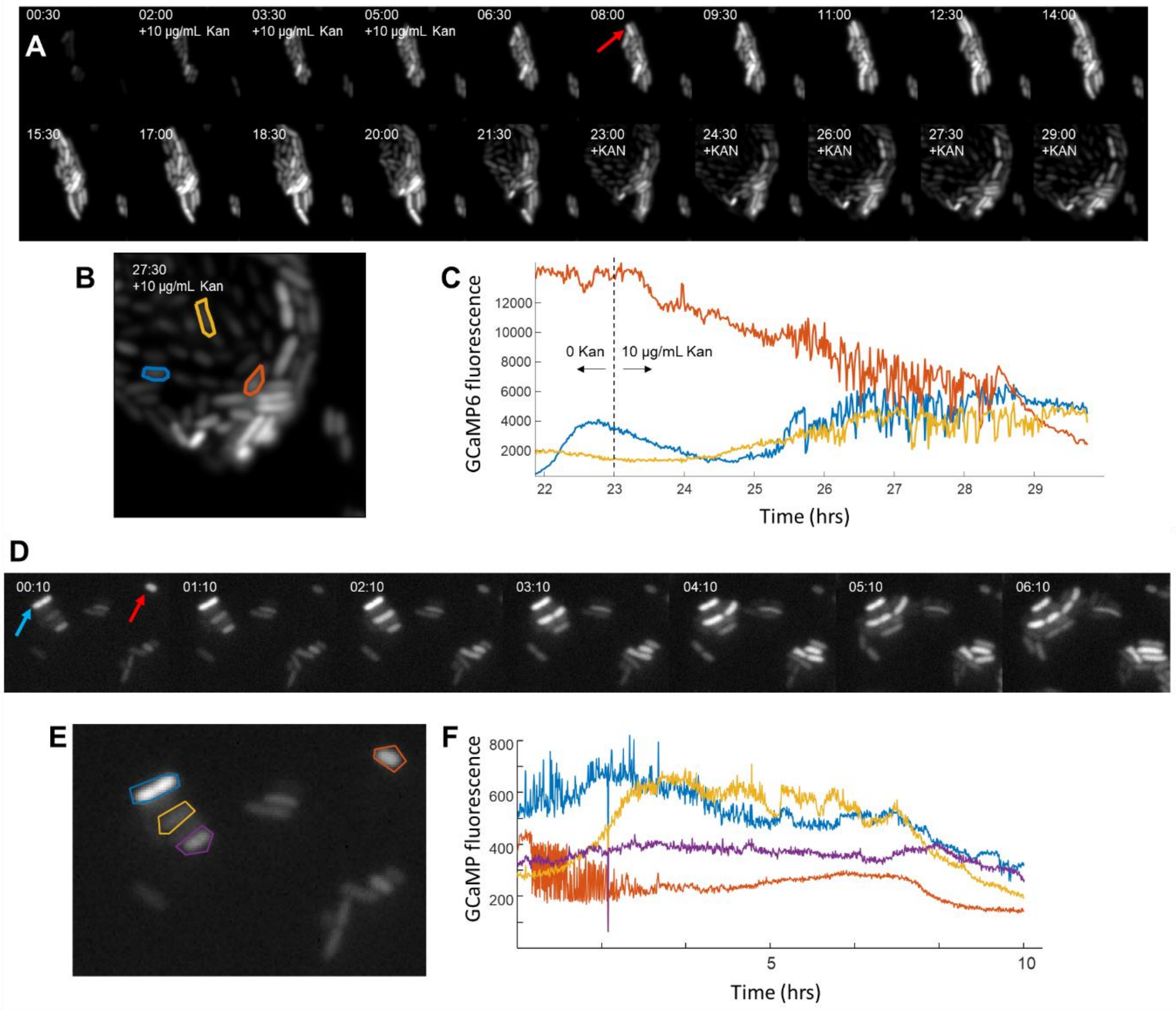


**Supplementary Figure 6. Related to Figure 3.**

(A-C) Cells that regrow after 1 treatment of kanamycin are not genetically resistant. (A) Strip chart of *E. coli* with 0 µg/mL kanamycin (t = 0-2 hours), 10 µg/mL kanamycin (t = 2-6 hours), 0 µg/mL kanamycin (t = 6-23 hours), and 10 µg/mL kanamycin (t = 23-30 hours). Time is shown at each frame in (HH:MM) format. The red arrow indicates a cell that regrows after the first kanamycin exposure. (B) Image from A at t = 27.5 hours. Individual cells were manually highlighted and shown with 3 different colors. Each cell has it's GCaMP6 time trace shown in (C) for just the time period of the second kanamycin addition. (C) GCaM6f fluorescence over time of individual cells in B. (D-F) Untreated cells that exhibit transients do not divide. (D) Strip chart of *E. coli* imaged via the GCaMP6 fluorescence under PMM without aminoglycoside treatment. The blue and red arrow mark cells that do not divide. The time is shown in (HH:MM) format. (E) Image taken after 10 minutes of imaging with individual cells segmented manually. Each color corresponds to the GCaMP6 fluorescent trace shown in (F). Any cell that exhibited transients in the untreated conditions did not divide within the movie, which ended with a fully overgrown field of view.

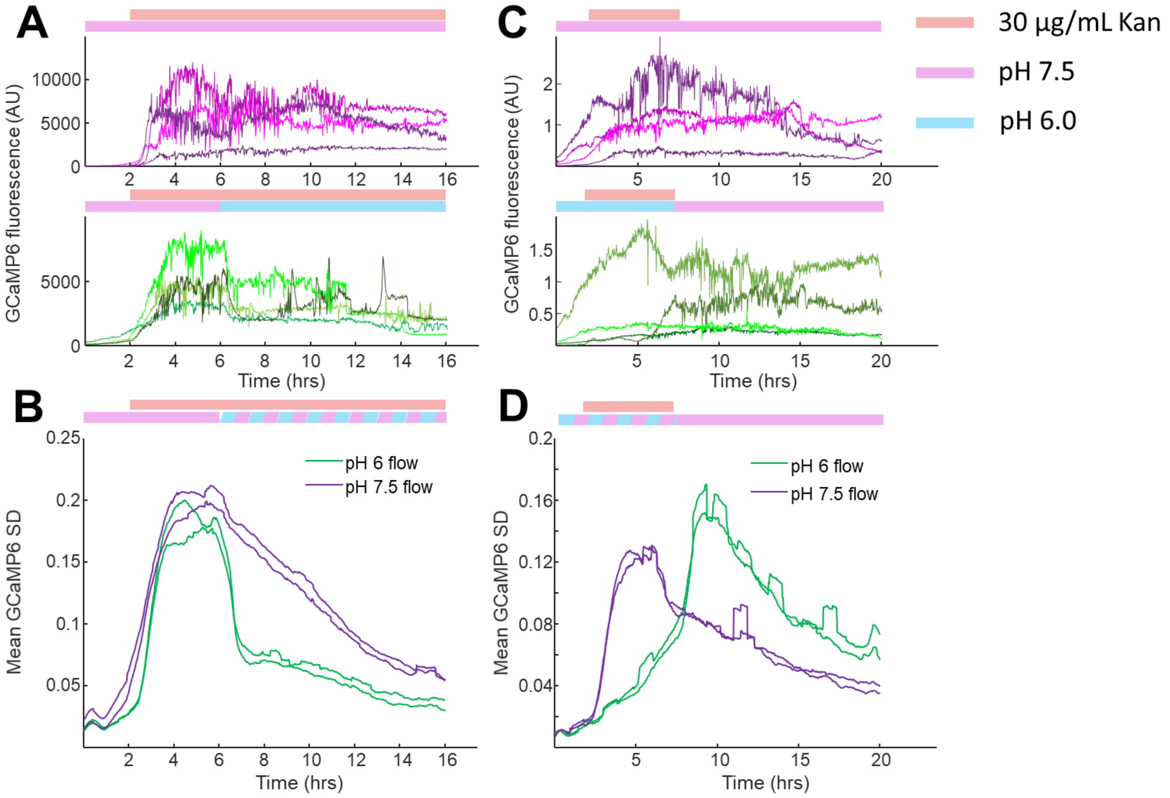


**Supplementary Figure 7. Related to Figure 4.**

(A-D) Catastrophic calcium transients require a membrane potential as indicated by low pH protection. (A) Random single cell traces of GCaMP6f intensity over time upon treatment with kanamycin (pink bar, top) at pH 7.5 (lilac bar, top), or with kanamycin and normal to low pH transition (lilac to blue bar, bottom). (B) Mean GCaMP6f standard deviation of biological replicates plotted over time. The population traces correspond to the mean of the single cell experiments in A, with kanamycin (purple, A-top, pH 7.5 only) or kanamycin +low pH treated cells (green, A-bottom, pH 7.5->6.0). (C) Random single cell traces of GCaMP6f intensity over time after kanamycin was flowed on top of the cells at 2 hours that were pretreated with PMM at pH 7.5 (top, lilac bar) or PMM at low pH (bottom, blue to lilac bar). Kanamycin and low pH were then flowed out of the chamber at 6 hours (pH was restored to 7.5, lilac bar). (D) Mean GCaMP6f standard deviation of biological replicates plotted over time. The population traces correspond to the mean of the single cell experiments in C, with kanamycin at constant pH 7.5 (purple, C-top, pH 7.5 only) or kanamycin +low pH transition treated cells (green, C-bottom, pH 6.0->7.5).

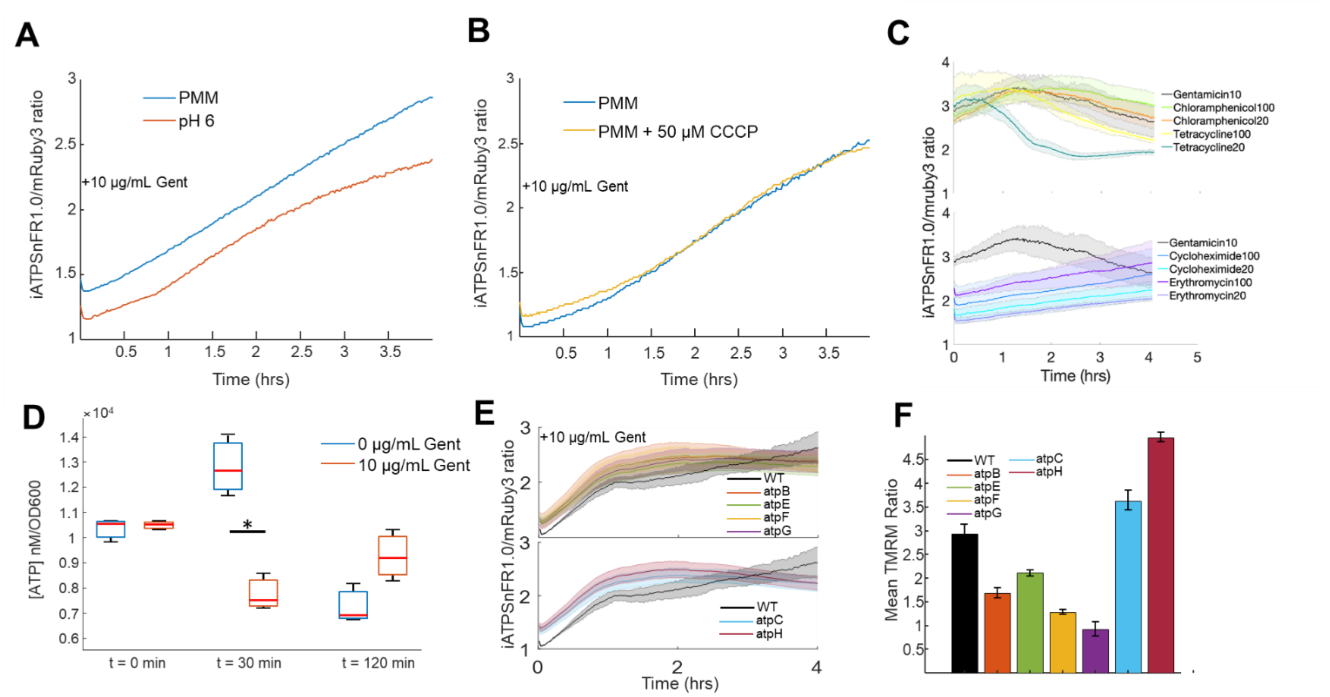


**Supplementary Figure 8. Related to Figure 5.**

(A,B) Low pH and CCCP maintain the rise in the iATPSnFR1.0/mRuby ratio upon gentamicin treatment when compared to controls. (A) ATP measured in PMM pH 7.5 (blue) or low pH (orange) upon treatment with 10 µg/mL gentamicin. Each line represents the average of 3 biological replicates. (B) ATP measured in PMM (blue) or PMM + 50 µM CCCP (yellow) upon treatment with 10 µg/mL gentamicin. Each line represents the average of 4 biological replicates. Even under protective conditions of CCCP or pH 6, 10 µg/mL gentamicin still induced a rise in ATP, but was not coupled with cell death. (C) Some bacteriostatic antibiotics have similar iATPSnFR1.0 ratios compared to gentamicin treated cells. (D) ATP concentration was quantified with a BacTiter Glo kit in the absence (blue boxplots) and presence (orange boxplots) of 10 µg/mL gentamicin at 0, 30, and 120 minutes. Simultaneously the optical density at 600 nm was read out, and the proportion of ATP to OD was taken to normalize the amount of ATP across the different time points and treatment conditions. The only significant difference between conditions is at 30 minutes with a p-value of 0.0036. (E,F) Compared to WT (black). (E) ATP concentration of 10 µg/mL gentamicin treated F1Fo-ATPase component knockouts. Each curve averages 4 biological replicates. (F) A bar plot of the ratio of the mean TMRM fluorescence from gentamicin treated (2 hours) to untreated cells measured by flow cytometry. Error bars are 95% confidence interval.

#### Movie captions

##### Movie S1.

Movie of *E. coli* expressing GCaMP6f-mScarlet upon treatment with 100 µg/mL kanamycin. The movie was taken using 488 nm excitation and a 40x air objective imaged onto an sCMOS camera. The movie was taken at a sampling rate of 1 image per minute for 16 hours. This movie has been corrected for uneven illumination, XY drift, and background as mentioned in the materials and methods. The time indicated represents HH:MM.

##### Movie S2.

Movie of *E. coli* expressing GCaMP6f-mScarlet with no kanamycin addition. The movie was taken using 488 nm excitation and a 40x air objective imaged onto an sCMOS camera. The movie was taken at a sampling rate of 1 image per minute for 16 hours. This movie has been corrected for uneven illumination, XY drift, and background as mentioned in the materials and methods. The time indicated represents HH:MM.

##### Movie S3.

Movie of *E. coli* expressing GCaMP6f-mScarlet switching the medium from PMM (0-5 hours), PMM + 10 µg/mL kanamycin (5-9 hours), PMM (9-35 hours). The movie was taken at a sampling rate of 1 image per minute for 29 hours. This movie has been corrected for uneven illumination, XY drift, and background as mentioned in the materials and methods. The time indicated represents HH:MM.

#### RESOURCES TABLE

| REAGENT or RESOURCE | SOURCE | IDENTIFIER |
|---|---|---|
| Bacterial and Virus Strains | | |
| *E. coli* K-12 BW25113 | Yale Coli Genetic Stock Center | CGSC#: 7636 |
| BW25113 ΔnuoA | Dharmacon Keio | OEC4987-213603796 |
| BW25113 ΔnuoB | Dharmacon Keio | OEC4987-213603795 |
| BW25113 ΔnuoH | Dharmacon Keio | OEC4987-213603791 |
| BW25113 ΔnuoI | Dharmacon Keio | OEC4987-213603790 |
| BW25113 ΔatpA | Dharmacon Keio | OEC4987-213606163 |
| BW25113 ΔatpB | Dharmacon Keio | OEC4987-213606167 |
| BW25113 ΔatpC | Dharmacon Keio | OEC4987-213605824 |
| BW25113 ΔatpD | Dharmacon Keio | OEC4987-213605825 |
| BW25113 ΔatpE | Dharmacon Keio | OEC4987-213606166 |
| BW25113 ΔatpF | Dharmacon Keio | OEC4987-213606165 |
| BW25113 ΔatpG | Dharmacon Keio | OEC4987-213607977 |
| BW25113 ΔatpH | Dharmacon Keio | OEC4987-213606164 |
| *E. coli* DK8 1100Δ(uncB-uncC)ilv::TnlO | Rubinstein lab (created in ref. 44) | DK8 |
| Chemicals, Peptides, and Recombinant Proteins | | |
| Glucose | Sigma | G7528-1KG |

| | | |
|---|---|---|
| M9 salts | Sigma | M6030 |
| MEM amino acid | Gibco | 11130-051 |
| Glutamate | Sigma | G1251 |
| Hydrochloric acid | Sigma | 320331 |
| Low melt agarose | VWR | 97064-134 |
| Sodium hydroxide | Sigma | 795429 |
| Kanamycin sulfate | Sigma | 60615-5G |
| Gentamicin sulfate | Sigma | 345814 |
| Apramycin | Sigma | A2024-1G |
| Streptomycin sulfate | Sigma | S9137 |
| Tobramycin | Sigma | 614005 |
| Trimethoprim | Sigma | T7883 |
| Cyclohexamide | Sigma | C7698 |
| Chloramphenicol | Sigma | C1919 |
| Erythromycin | Sigma | E5389 |
| Potassium chloride | Sigma | P9333 |
| Magnesium chloride | Sigma | 63068 |
| CCCP | Sigma | C2759 |
| Oxyrase for broth | Sigma | SAE0013 |
| Texas Red-X, Succinimidyl Ester | Thermo-Fischer | T20175 |
| N,N-Dimethylformamide | Sigma | 227056 |
| Glutathione | Sigma | G4251 |
| Ascorbic acid | Sigma | A7506 |
| Propidium iodide | Life-tech | P3566 |
| Tetramethylrhodamine, methyl ester | Molecular Probes | T668 |
| DL-Dithiothreitol | Sigma | D9779 |
| Lysozyme from chicken egg white | Sigma | 62971-10G-F |
| Magnesium chloride hexahydrate | Sigma | 63068-250G |
| Sucrose | Sigma | 84097-1KG |
| Water, Sterile. WFI Quality | Sigma | 4.86505.1000 |
| Sodium Deoxycholate | Sigma | 30970-25G |
| Ammonium chloride | Sigma | 09718-250G |
| Recombinant DNA | | |
| Plasmid: pKL09-GCaMP6f-mScarlet bb118 | This paper | Addgene #pending |
| Plasmid: pKL10-mRuby-iATPSnFR1.0 bb118 | This paper | Addgene # pending |
| Plasmid: pKL12-GCaMP pHuji bb118 | This paper | Addgene # pending |
| Plasmid: pKL11-GCaMP6f bb100 | This paper | Addgene # pending |
| Plasmid: pKL13-MgtC bb118 | This paper | Addgene # pending |
| Software and Algorithms | | |
| MATLAB | https://www.mathworks.com/products/matlab.html | RRID:SCR_001622 |
| NIS Elements | https://www.microscope.healthcare.nikon.com/products/software/nis-elements | RRID:SCR_002776 |

**Material and Methods**

LEAD CONTACT AND MATERIALS AVAILABILITY

Plasmids generated in this study are available on Addgene. Knockout strains from the Keio collection are available through Dharmacon due to an MTA. Reagents, data and analysis code are available upon request, and will be fulfilled by the Lead Contact, Joel Kralj.

EXPERIMENTAL MODEL AND SUBJECT DETAILS

*E. coli Strains*

*E. coli* strain BW25113 was acquired from the Yale Coli Genetic Stock Center and was used as the control, except experiments where specifically noted. Knockout strains were acquired from the Keio collection purchased from Dharmacon (#OEC4988). *E. coli* strain DK8 1100Δ(uncB-uncC)ilv::TnlO, which is deficient of the F-ATPase was a generous gift from the Rubinstein lab.

*Cell growth*

Strains were grown in LB with antibiotics dependent on growth conditions. For GCaMPmScarlet expressing cells, clones transformed with the plasmid were grown overnight with carbenicillin (100 μg/mL). Carbenicillin was used for overnight cultures to maintain the plasmid, but was not present for any experiments. For knockout strains from the keio collection kanamycin (50 μg/mL) was also added to any overnight cultures. Strain DK8 was grown overnight in the presence of tetracycline (30 μg/mL). Glycerol stocks were streaked onto plates bearing the appropriate antibiotics, and individual colonies were picked and grown in 5 mL culture tubes, or in 24 well plates, or in 50 mL Erlenmeyer flasks. All cells were grown overnight at 37 ̊C with shaking between 150-200 rpm with the appropriate antibiotic if required for plasmid or strain selection. Knockouts from the Keio collection were plated on LB plates with kanamycin and carbenicillin to ensure maintenance of the knockout cassette, but overnight liquid cultures that were to be used for imaging were grown only in the presence of carbenicillin to avoid any potential effects of protein translation inhibition on sensor expression.

METHOD DETAILS

*Plasmids*

Expression of GCaMP6f-mScarlet was carried out with a constitutive promoter (118, iGem biobrick) assembled in an ampicillin resistant plasmid similar to earlier work(*1*). The mScarlet amino acid sequence was taken from the original publication(*2*) and purchased as a gBlock (IDT). The plasmid was double digested with Pme1/Nco1 and assembled using Gibson assembly. The mRuby-iATPSnFR1.0 construct was created by obtaining the amino acid sequence directly from the publication(*3*) and codon optimizing it in a single gBlock ordered from IDT, then Gibson cloned into the same constitutive promotor backbone as GCaMP6f-mScarlet. Expression of these constructs was carried out in the 118 plasmid. Expression of GCaMP6f alone was carried out using a similar constitutive promoter (100, iGem biobrick) in the same backbone. The mgtC over expression plasmid was created by obtaining the amino acid sequence directly from salmonella on a gBlock, and Gibson cloned into the 118 biobrick backbone used above. GCaMP6f tethered to pHuji was purchased on a gBlock and Gibson cloned into the same constitutive biobrick 118 promoter used previously. All plasmids and sequences have been deposited on Addgene. All plasmids were transformed into their respective genetic background strain using Transfer Storage Solution transformation protocol.

*Imaging media and fluorescent dyes*

Unless otherwise noted, all imaging experiments were conducted in PMM at pH 7.5. The PMM recipe used is: 1x M9 salts (Sigma), 0.2% glucose (Sigma), 0.2 mM $MgSO_4$, 10 µM $CaCl_2$, 1x MEM amino acids (Gibco). Experiments were conducted at pH 7.5 unless otherwise noted in the text, and NaOH or HCl was used to change the pH to the final value. Given the critical importance of pH in aminoglycoside response, all PMM media with additional chemicals was pH adjusted to 7.5 before imaging. At more basic pH, and higher concentrations of Mg, precipitate forms in this media over time. For oxygen free microscopy experiments, Oxyrase for Broth was added to the media pads during the pre-imaging incubation time to 10% v:v, then sealed to have oxygen removed.

Propidium iodide (Life Tech) was dissolved in water in a stock concentration, and added to a final concentration of 3 µg/mL. PI was imaged with a 561 nm laser in a flow experiment, and was added at the same time as 30 µg/mL kanamycin.

TMRM (ThermoFischer) was dissolved in DMSO in a 1 mM stock solution, and diluted in PMM to 8 µM, then added to a final concentration of 200 nM to cell suspensions. TMRM was measured as described below in flow cytometry.

*Preparing cells for imaging*

All imaging of cells took place under agarose pads which were composed of PMM at the appropriate pH and 2% low melt agarose.

For experiments using flow, the agarose was melted in PMM buffer and cast between 2 pieces of glass covering a silicone mold. The silicone was 3/16" as the final thickness, and was cut by hand. The pads were diced into small squares using an exacto knife to fit into the flow chambers (~2 mm x 2 mm). Cells from an overnight culture were placed directly on to the agarose pad (1.0 µL) and left for ~5 minutes. The agarose pads were then placed with the cells down onto a 24 mm x 50 mm glass coverslip (thickness 1.5) with a silicone flow chamber. The apparatus was then sealed with a custom glass slide with holes drilled to enable flow.

Experiments involving drug titrations or knockouts were prepared onto 96 well glass bottom dishes (Brooks Automation, MGB096-1-2-LG-L). A custom 96 well mold was created and 3D printed using a commercial service. The mold was designed to hold a volume of 200 µL per well (Shapeways), with a separate piece designed to press the agarose pads into the coverslip in 8, 12, or 96 well format. The 3D printed pieces are available at the Kralj Lab store on Shapeways (https://www.shapeways.com/shops/kraljlab) and the .stl files are available to researchers upon request. The bottom of the agarose mold was sealed with a 4" x 6" piece of glass (McMaster Carr), and liquid agarose was added to the desired wells. A second piece of glass (3" x 5") was used to seal both sides, and the agarose was left to cast for > 1 hour. The glass piece was then removed, and cells were added to each pad individually (2 µL) and left for 10 minutes for the liquid to absorb into the agarose. The cells were then pressed out into the 96 well plate using the custom 3D printed press. For all experiments, cells were left in the pad for ~1 hour before imaging. Any chemical treatments were then added to the top of the pad. A 5 µL drop of a solution at 40x final concentration was added on to the pad and left to diffuse throughout. In house measurements with a small fluorescent dye showed compounds diffuse to the glass in ~5 minutes.

*Imaging*

Flow experiments were conducted using a Nikon TiE base with perfect focus, running Elements software, with a custom laser illumination with high angle illumination. A 488 nm (Obis 150 LX, Coherent) or 561 nm (Obis 50 LS) were combined, expanded, and focused onto the back aperture to create a widefield illumination. A mirror located 1f away from the widefield lens was used to control the illumination angle. Imaging took place with a 100x NA 1.45 objective with intensities (at the sample) of 130 $mW/cm^2$ 488 nM and 1050 $mW/cm^2$ 561 nm light. A quad band emission filter (Semrock) was used for reflecting the illumination light, and no emission filter was needed. The light was imaged onto an Andor EMCCD (iXon 888 Ultra) using an exposure time of 200 ms. Images were acquired sequentially (561nm, then 488 nm) once per minute over the entire experiment (6 – 48 hours). These illuminations showed no evidence of phototoxicity compared to unilluminated cells as measured by growth rate.

Flow was controlled with two identical syringe pumps (Harvard Apparatus). Flow rates were set to 20 µL/minute which was sufficient to fully exchange the medium in the chamber within 2.5 minutes. Each syringe pump was loaded with the appropriate medium and was programmed to turn on or off at the desired time. A typical experiment involved 2 hours of PMM alone, followed by switching to PMM+Kan using the second pump. Tubing from multiple syringes was connected with a T-connector with a dead volume of ~20 µL. At all times during flow cell experiments, the specified media was flowed through the chamber.

Imaging 96 well glass bottom plates took place using a Nikon Ti2 inverted microscope running the Elements software package. Fluorescent excitation was achieved with a Spectra-X LED source (Lumencor). A 40x, NA 0.95 air objective was used to both illuminate and image the cells onto 2- Flash 4 v2 sCMOS cameras (Hamamatsu) using a custom splitter to image 2 colors simultaneously (Thorlabs). Illumination was achieved by simultaneous excitation with 470/26 and 554/20 band pass LED illumination for a 200 ms exposure. Measured light intensities at the sample were 330 $mW/cm^2$ (470 nm) and 2050 $mW/cm^2$ (554 nm). Typical sampling rates were 1 frame per minute, unless noted in the text.

*CFU measurements*

CFUs were measured by plating treated cells onto LB-agarose without antibiotic and counting growing colonies. CFU measurements were conducted trying to mimic the experiments performed via microscopy. Briefly, cells were grown overnight in LB and diluted 1:20 in 5 mL PMM. These cultures were grown at room temperature and shaking for 2 hours (t = 0) followed by the addition of antibiotic. At each time point, the culture was removed from the shaker, and 100 µL was removed. A 10x series dilution was then conducted by removing 20 µL and adding to 180 µL LB alone in a 96 well plate. The 10-fold dilution was performed 7 times, leading to the original concentration to a dilution of $10^7$. From each of the 10x dilution series, 3 µL was plated onto an LB agar pad and left to dry (1 colony = 333 cells/mL, lower end of our dynamic range).  After an entire experiment (typically 5 hours), the agar was placed into an incubator and grown overnight.  Colonies were then manually counted the next morning.

*Cell cytometry*
A 5 ml of PMM media was seeded with 50 µL of overnight BW25113 cells, or the respective knockout strain tested. When the cells reached ~0.4 OD, 100 ug/ml kanamycin, 10 ug/ml gentamicin, or PMM alone was added. After 30 minutes of antibiotic or mock treatment, TMRM was added to the suspension at a final concentration of 0.2 mM. Two hours later, 1 mL of cell suspension was transferred to a 5 mL Falcon polystyrene round-bottom tube. Cells were quantified for their TMRM incorporation by counting 100,000 events per condition using a BDFACSCellesta Flow Cytometer with the following Voltage settings: FSC at 700, SSC at 350, with 561 nm laser D585/15 at 500, C610/20 at 500 and B670/30 at 481. Emission for each event was collected at the 585/15 nm wavelengths.

*Bactiter glo ATP analyses*
ATP per optical density unit was quantified using Promega's BacTiter-Glo kit coupled with a BioTek Synergy plate reader. BacTiter-Glo reagents and standards were prepared as described in the manual. Briefly, exponentially growing cultures of *E. coli* were treated according to the experimental parameters for the times indicated. When the time of treatment was reached 100µL of culture, blank, or standard, was added to a black walled clear bottom 96-well plate. This was done in technical triplicate for each condition, blank, or standard, which had at least three biological replicates. Once the plate was prepared 100 µL of BacTiter-Glo Reagent was added to each well, and shaken in an orbital shaker for 1 minute at room temperature, then left on the benchtop for 5 minutes. Luminescence was recorded using the BioTek Synergy plate reader, set to auto scaling and 1 second integration time. Simultaneously with BacTiter-Glo plate preparation, an optical density plate was created with the same cultures, and the absorbance of the culture was read on the same plate reader at 600 nm. ATP per OD unit was calculated by the average of the three biological replicates (which were averaged from the technical replicates), which were then divided by the obtained OD values.

*Polysome analyses*
Sucrose gradients were prepared in Beckman Coulter Ultra-Clear™ Tubes (14x89mm) Reorder No. 344059. Media recipes and protocol is from (*5*).  Roughly 6 ml of 10% sucrose was layered on the bottom of the tube, then a large needle was used to add 40% sucrose below the 10% layer up to a 6 ml marker on the outside of the tube. If a clear meniscus between the two layers was not visible the tube was discarded. Tubes were placed in a MagnaBase™ tube holder (sku B105-914A-I/R), and short caps were placed on top to eliminate all air from the tube. The tube holder was then placed on the gradient maker.  A 10-40% gradient was then established using a BIOCOMP Gradient Station ip gradient maker with the following settings: Short cap, Sucrose, 10-40%wv, 81˚, 1:48 min:sec. Caps were then removed, and gradient tubes were stored no longer than 1 hour at 4˚C until lysate supernatant was prepared.

Ribosomes and ribosomal subunits were characterized using a slightly adapted protocol, due to differences in available equipment, from (*5*).  Briefly 50 mL cultures were grown to ~0.35-0.45 OD.  Antibiotic, or a mock treatment was then added, and these cultures were allowed to grow for another 1 (LB, Fig 5A,B) or 1.5 (PMM, Fig 5C) hours.  Optical Density was taken at time of collection, when 37.5 ml of culture was then transferred to Nalgene® Oak Ridge Centrifuge Tubes (Cat. 3119-0050) on ice. Cells were then pelleted in a chilled Sorvall SA-600 rotor in a Sorvall RC 5C Plus Centrifuge at 10000 rpm for 5 minutes at 4˚C. Culture media was decanted and aspirated. Cell pellets were then resuspended in 500 µL lysis buffer (750 µL for anaerobic conditions), and flash frozen in liquid nitrogen. Frozen suspensions were thawed in a 5-10˚C water bath, then flash frozen again, and either stored at -80˚C or thawed in the same manner and treated as follows. Lysis was completed by adding 15 µL of 10% sodium deoxycholate to freeze-fractured pellet resuspensions and mixed by inversion. Lysate was then separated by centrifugation at 4˚C at 10,000 rpm in a chilled Eppendorf FA45-30-11 rotor in an Eppendorf 5804R Centrifuge for 10 minutes at 4˚C. Lysate supernatant was collected in chilled microfuge tubes. Then 300 µL of the 10-40% gradient was removed from the top of the sucrose gradient columns and replaced with 300 µL of lysate supernatant.

Loaded gradient columns were placed in Beckman SW-41 swinging buckets and balanced to within 0.01 g of each other using the 10% sucrose solution. Loaded sucrose gradient buckets were then centrifuged using the SW-41 rotor in an LM-8 Ultracentrifuge in 4˚C at 35,000 rpm for 3 hours. Sucrose gradient columns were then removed, and fractions were then collected using the following series of machines. A BIOCOMP Gradient Station IP with settings Distance 80.00 mm, Speed 0.3 mm/sec was tethered to a BIORAD Model 2110 Fraction Collector with the following settings: 6 drops/fraction. As fractions were collected the absorbance at 254 nm was collected from the fractions using a BIORAD Econo UV Monitor set to range 1.0 (AUFS) tethered to computer running the BIOCOMP Gradient Profiler 2.0 software. Data files for each gradient run were saved as .csv files and later analyzed in Matlab using custom scripts to integrate peaks with the trapz.m function.

Due to the nature of collection with these devices, often the beginning of the non-ribosomal RNA peaks was missed, capturing the absorbance as the non-ribosomal RNA ran through the detector midway through the peak. In all conditions tested, non-ribosomal RNA, 30S, 50S, and 70S peaks were detected. To simplify comparisons between conditions, polysomes beyond the 70S peak were ignored in the (30S+50S)/70S ratio measurements. Note that because of the nature of these experiments, different total quantities cell lysate, and therefore of total RNA, are loaded into the sucrose gradients columns. Due to this reality, comparing the 254nm absorbance quantities between samples is unreasonable, however comparing the ratio of the ribosome peaks should be total-RNA agnostic.

#### QUANTIFICATION AND STATISTICAL ANALYSIS

*Image processing*

Data was stored as .ND2 files which contain the 16 bit images and the associated metadata. The BioFormats Matlab package was used to access data in the .ND2 format. All data analysis was performed using custom scripts in Matlab (available upon request).

Image processing followed the general scheme of (1) estimating the illumination profile for all experiments on a given day, (2) correcting the uneven illumination for each movie, (3) registering drift and jitter in XY, (4) subtracting an estimated background, (5) segmenting cells using a Hessian algorithm, (6) extracting time traces for individual cells, (7) processing each time trace for the onset and amplitude of calcium transients.

1. Estimating the illumination profile: For a given day, every movie was averaged across time, and opened using a morphological operator and blurred using a 2D Gaussian filter. Each of these experimental images were then averaged together to give an estimate of the uneven illumination. These images were smooth across the entire field of view, and varied by ~50% across the entire image.

2. Correcting uneven illumination. Each individual movie was then loaded into memory sequentially. Each frame of the movie was converted to a double, and then divided by the uneven illumination. This image was then multiplied by the average value of the movie and converted back into a uint16 to maintain consistent intensity values. Each frame was then reassembled into an illumination corrected movie.

3. Registering drift and jitter in XY: Each frame was aligned to the previous frame using a convolution of the 2D Fourier transform. Each sequential image was first estimated by applying the XY warping from the previous frame. Then, the 2DFT was taken for each image, and multiplied to the previous frame. The optimal updated XY position was then calculated and applied.

4. Subtracting the estimated background: The background was estimated for each frame individually using a morphological operator. A disk structured element with radius 9 µm was blurred with a Gaussian filter. This background estimation was then subtracted from the original image. To protect against potential negative values, the minimum of the entire movie was set to 50 counts.

5. Segmenting cells using a Hessian algorithm: To segment cells, first the foreground was estimated using Otsu's method from the background subtracted image. The Hessian was then calculated on the background subtracted image, and then elementwise multiplied to a logical image of the foreground. Otsu's method was again used on this modified Hessian image to identify individual cells. Hard limits were set to remove potential noise that did not fit given criteria for size or minimum intensity. We found that first increasing the size of the image using a spline interpolation gave superior segmentation results. Using this method, not all cells were identified within a microcolony, though we estimate that it can identify ~96% of the cells accurately.

6. Extracting time traces for individual cells: From a given identified cell, for each time point in the movie, we extracted the mean intensity using the Matlab command, regionprops. The mean intensity for both the GCaMP6f and the mScarlet were extracted using this method, or any other fluorophore the cells expressed.

7. Processing each time trace for the onset and amplitude of transients: For each time trace, the moving median over 45 minutes was divided to remove the slow baseline trends. A standard deviation was calculated from the timepoints before aminoglycoside addition, and a cell was defined as blinking if it had transients that lasted > 10 minutes that were > 7x the pre-treatment

standard deviation. From each normalized time trace, the moving standard deviation was also calculated using a 30-minute sliding window. Within a given FOV, the entire population moving standard deviations was averaged, providing the average standard deviation trace shown in the figures.

During flow experiments, single frames were sometimes contaminated by bubbles that dramatically changed the contrast. To remove these features, we took the average of all extracted cells. If the differential of any single frame was initially lower, then higher than 5x the standard deviation of the whole movie, a single frame was removed. This preprocessing removed spurious catastrophic blinks that appeared to occur in every cell at the same instant.

*Cytometry analysis*
Analysis of the cytometry data was achieved by fitting the data to a 1D Gaussian distribution and calculating the mean and 95% confidence interval for each of these fits for each strain tested. These values were then taken as a ratio of the gentamycin treated cells relative to the vehicle treated cells.

Assuming TMRM partitions according to Boltzmann's law:

$$\frac{C_{in}}{C_{out}} = e^{\frac{-q}{kT}V},$$

where $C_{in}$ and $C_{out}$ are the concentrations of the dye in and out of the cell, q is the ionic charge, k is the Boltzmann constant, and T is the temperature in kelvin. Comparing two different conditions ($V_{kan}$ and $V_{PMM}$), we can solve for the treated condition to yield:

$$V_{Kan} = V_{PMM} - 28mV * \ln\left(\frac{C_{in,Kan}}{C_{in,PMM}}\right)$$

if we assume the concentration of dye out of the cell is the same in both conditions. Given a large reservoir relative to the cytoplasmic volume of the cells, this is a reasonable estimate.

*Significance testing*
Significant differences across populations of individual cells were tested using the unpaired t-test with unequal variance. For cytometry experiments, we used the 95% confidence interval (CI) to a single Gaussian fit. All significance testing has been summarized in Table 1 for simplification of figure representation.

DATA AND CODE AVAILABILITY
The datasets and code supporting the current study have not been deposited in a public repository because the of the size of the files but are available from the corresponding author on request.
